# The role of dopamine-sensitive motor cortical circuits in the development and execution of skilled forelimb movements

**DOI:** 10.1101/2025.06.20.660525

**Authors:** Martyna Gorkowska-Nosal, Gniewosz Drwiega, Lukasz Szumiec, Jan Rodriguez Parkitna, Przemyslaw E. Cieslak

**Affiliations:** Department of Neurophysiology and Chronobiology, Institute of Zoology and Biomedical Research, Faculty of Biology, Jagiellonian University, 30-387 Krakow, Poland; Doctoral School of Exact and Natural Sciences, Jagiellonian University, 30-348 Krakow, Poland; Department of Molecular Neuropharmacology, Maj Institute of Pharmacology Polish Academy of Sciences, 31-343 Krakow, Poland; Present address: Department of Computational Biophysics and Bioinformatics, Faculty of Biochemistry, Biophysics and Biotechnology, Jagiellonian University, 30-387 Krakow, Poland

**Keywords:** motor cortex, skill learning, dopamine, fiber photometry, head-fixed behavior

## Abstract

Dopamine signalling in the motor cortex is crucial for motor skill learning. Here we resolve the spatiotemporal dopamine dynamics and the activity of local dopaminoceptive circuits during the formation and execution of motor skills. We trained head-fixed mice to perform skilled forelimb movements with a joystick to collect water rewards, while simultaneously monitoring dopamine release and calcium dynamics in the forelimb area of motor cortex. We found that dopamine release events and calcium transients were temporally linked to joystick movements and reward consumption. Dopamine dynamics and population level activity of dopamine-receptive neurons scaled with the vigor of forelimb movements and tracked the relationship between actions and their consequences. Optogenetic photoinhibition of cortical dopaminoceptive circuits reduced the number of rewarded joystick movements. Our findings show how phasic dopamine signals in the motor cortex facilitate reinforcement motor learning of skilled behavior.

## Introduction

The acquisition of new motor skills and the adaptation of motor behavior to changing environmental demands are essential for survival. A distributed neural network mediates the learning and performance of motor skills, with the primary motor cortex (M1) acting as a central hub that transmits output cortical motor commands to downstream motor centers, including the basal ganglia, motor thalamus, brainstem, and spinal cord^1–4^. The M1 is innervated by dopaminergic fibers, with preferential innervation in the M1 forelimb area^5–10^. There is growing evidence that dopamine (DA) transmission is crucial for motor skill learning, mediates structural and cellular synaptic plasticity in the M1, and modulates reach kinematics^6,11–15^. Nevertheless, the spatiotemporal DA dynamics in the M1 during the formation and execution of skilled forelimb movements have not yet been investigated.

DA action in M1 is mediated by DA receptors D1 and D2, encoded by the *Drd1a* and *Drd2* genes, respectively. Application of specific D1 and D2 antagonists into the M1 forelimb area suppresses synaptic plasticity and motor skill acquisition^11,12,16^. Moreover, activation of D1 and D2 receptors has been shown to modulate the excitability of M1 pyramidal and GABAergic interneurons^10,17,18^. Nevertheless, still little is known about M1 populations of dopaminoceptive neurons. Recently, a few studies have employed transgenic mouse strains, with *Drd1a* and *Drd2* gene promoters driving the expression of Cre recombinase and fluorescent proteins, to selectively target populations of D1 receptor-positive (D1+) and D2 receptor-positive (D2+) cells in the mouse M1^17–19^. These studies showed that D1+ and D2+ populations are non-overlapping (expressing either D1 or D2 receptors), diverse (containing pyramidal neurons and GABAergic interneurons), and have a preferential laminar distribution (D1+ neurons are mainly found in the deep layers, whereas D2+ primarily found in the superficial layers). However, the activity patterns of these populations during movement execution and their functional contribution to motor performance remains to be determined.

We have developed a forelimb-specific task in head-fixed mice, in which animals make skilled forelimb movements using a joystick to acquire water reward^20–22^. We used fiber photometry to track DA dynamics and population-level calcium (Ca^2+^) activity in the M1 of D1Cre and D2Cre mice, during the development of skilled behavior and following subsequent changes in reward threshold and contingency. Furthermore, we used optogenetic photoinhibition to investigate the functional contribution of the M1 D1+ and D2+ populations to motor performance and employed retrograde tracings to determine their long-range connections. We provide the first comprehensive assessment of the role that DA-sensitive motor cortical circuits play in the formation and execution of skilled forelimb movements.

## Results

### Simultaneous recording of DA dynamics and population level Ca^2+^ activity in the M1 forelimb area

We recently identified two populations of DA receptor-expressing neurons with a layer-specific distribution in the forelimb area of the mouse M1^19^. We showed that D1 receptor-positive (D1+) neurons are primarily found in the deep layers, whereas D2 receptor-positive (D2+) cells are distributed in the superficial layers. Here, we wanted to determine DA release dynamics across the deep and superficial layers of the M1, and record cell type-specific Ca^2+^ activity in the previously identified populations of DA-sensitive neurons in awake, behaving animals. We injected an adeno-associated virus (AAV) expressing a high-affinity green fluorescent DA sensor (GRAB_DA2h_)^23^ and Cre-dependent red fluorescent Ca^2+^ indicator (jRGECO1a)^24^ into the M1 forelimb area of D1Cre and D2Cre transgenic mice^25,26^, expressing Cre recombinase in dopaminoceptive neurons (**Figure 1A**). In Cre-littermates, only the DA sensor was expressed. AAV injections were made in layer 5 (L5) of D1Cre mice and layer 2/3 (L2/3) of D2Cre animals, with optical fibers positioned above the injection site in the right hemisphere (**Figure 1B**). Using CMOS fiber photometry system^27^, we simultaneously monitored GRAB_DA2h_ and jRGECO1a signals in head-fixed mice trained to make skilled, bimanual forelimb movements with a joystick to acquire a delayed water reward (**Figure 1C)**. We were able to detect high-frequency Ca^2+^ transients in both populations of D1+ and D2+ neurons, and corresponding DA release events in L5 and L2/3. Throughout the recording sessions, the frequency of Ca^2+^ transients remained constant, while DA release events gradually increased (**Figures 1D and 1E**). When we compared fiber photometry data with video recordings of mice performing the task, we observed that neuronal activity in the M1 was specifically linked to forelimb motions, ranging from small subthreshold joystick movements or paw placement adjustments to large forelimb motions including joystick manipulations, spout reaching, or grooming (**Figures 1D and 1E**). Overall, we verified that by using fiber photometry we can record DA dynamics and population Ca^2+^ activity in DA-sensitive neurons in the M1.

**Figure 1.**
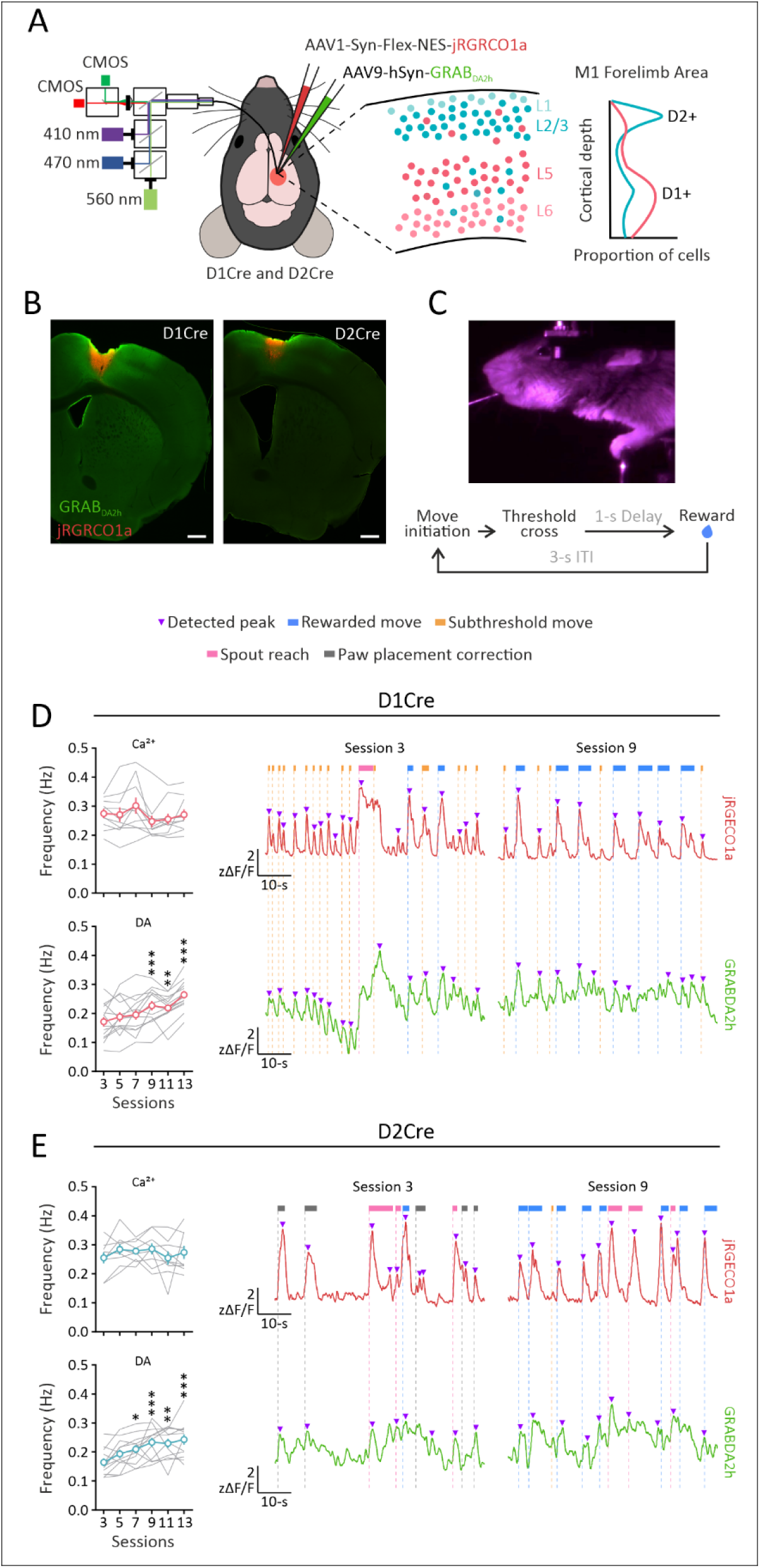
Simultaneous recording of DA dynamics and population level Ca^2+^ activity in the M1 forelimb area. (A) Diagram of the fiber photometry recording setup and schematic of laminar distribution of D1+ and D2+ neurons in the M1 forelimb area (based on ^19^). (B) Coronal sections obtained from a D1Cre and D2Cre mice showing expression of GRAB_DA2h_ and jRGECO1a. Scale bars, 500 µm. (C) Head-fixed mouse performing joystick task and the trial structure schematic. (D) Fiber photometry recordings in D1Cre mice. (Left) Frequency of the D1+ population Ca^2+^ transients (n = 10) and DA release events (n = 14). (Right) Z scored jRGECO1a and GRAB_DA2h_ traces recorded simultaneously in L5 of D1Cre mouse, obtained from sessions 3 and 9. See also **Videos S1-S4**. (E) Fiber photometry recordings in D2Cre mice. (Left) Frequency of the D2+ population Ca^2+^ transients (n = 9) and DA release events (n = 12). (Right) Z scored jRGECO1a and GRAB_DA2h_ traces simultaneously recorded in L2/3 of D2Cre mouse, obtained from sessions 3 and 9. See also **Videos S5-S8**. (D and E) Data are represented as mean ± SEM. Repeated-measure one-way ANOVA; multiple comparisons: S3 mean vs. every other mean; Bonferroni’s post hoc test; \*\*\**p* < 0.001, \*\**p* < 0.01, \**p* < 0.05.

### Head-fixed mice learn to perform skilled forelimb movements to obtain reward

As training progressed, task-unrelated forelimb motions were gradually replaced by forelimb movements involving the use of a joystick, and mice of both genotypes demonstrated a significant increase in the number of rewarded trials (**Figures 2A and 2D**). To further investigate the development of this skilled behavior, we compared the movement kinematic parameters between the ‘early’ and ‘late’ learning sessions. For the ‘early’ session, we selected a day on which the animal made at least 30 rewarded joystick movements (median number of sessions: D1Cre – 4.5; D2Cre – 4; median number of rewarded trials: D1Cre – 75; D2Cre – 76). The last two days were used for the ‘late’ session comparison (median number of sessions: D1Cre – 14; D2Cre – 14; median number of rewarded trials: D1Cre – 163; D2Cre – 174). We found that the mice gradually produced more vigorous movements, as the amplitude and velocity increased significantly across the learning days (**Figures 2B and 2E**). Since reward was elicited by joystick movements in any direction, we saw a considerable degree of trial-to-trial variability in movement trajectories (**Figures 2C and 2F**). Even though the movements varied in direction and amplitude, the mice developed a tendency to perform structured push-pull motions along the body axis during training. Overall, we demonstrated that head-fixed animals developed goal-directed behavior and used skilled forelimb movements to obtain delayed rewards.

**Figure 2.**
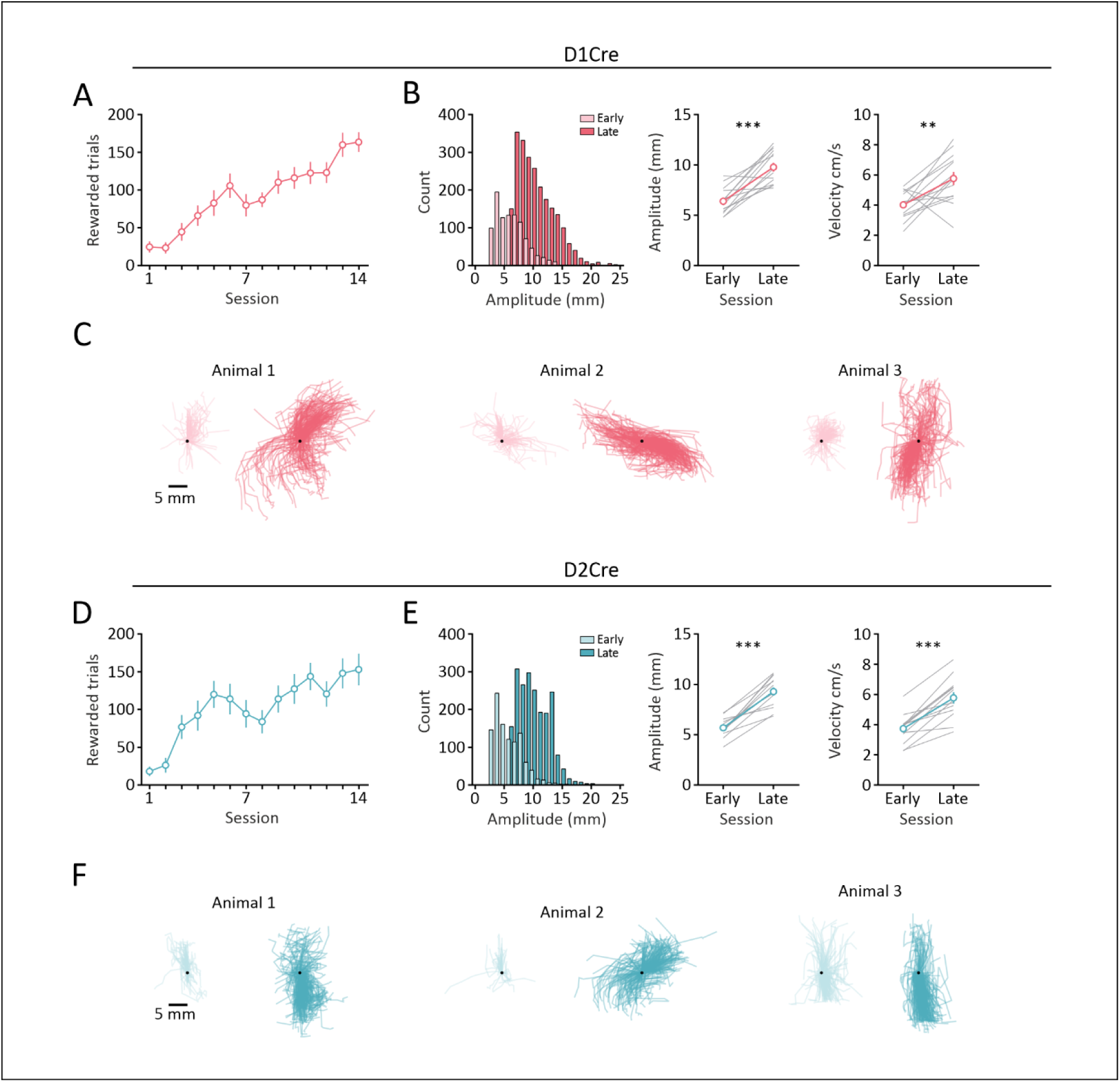
Head-fixed mice learn to perform skilled forelimb movements to obtain reward. (A) The number of rewarded joystick movements during training in D1Cre mice (n = 14). (B) Movement kinematic parameters obtained from the ‘early’ and ‘late’ sessions in D1Cre mice. (Left to right) Distribution of movement amplitudes; movement amplitude; joystick velocity at the time of threshold crossing (n = 14). (C) Movement trajectories of representative D1Cre mice aligned to the initial joystick position (black dot). (D) The number of rewarded joystick movements during training in D2Cre mice (n = 13). (E) Movement kinematic parameters obtained from the ‘early’ and ‘late’ sessions in D2Cre mice. (Left to right) Distribution of movement amplitudes; movement amplitude; joystick velocity at the time of threshold crossing (n = 13). (F) Movement trajectories of representative D2Cre mice aligned to the initial joystick position (black dot). (A, B, D and E) Data are represented as mean ± SEM. (B and E) Paired t-test; \*\*\**p* < 0.001, \*\**p* < 0.01.

### DA dynamics in the M1 encode execution of skilled forelimb movement and reward consumption

Next, we mapped DA and Ca^2+^ activity in the M1 and licking patterns following joystick movement and reward delivery, throughout the same ‘early’ and ‘late’ sessions (**Figure 3**). Data were aligned to the threshold crossing (**T**: 0 s), with the preceding 2 s serving as the baseline (**BL**: -2 to 0 s), and analyzed at 1 s intervals. As demonstrated in D1Cre and D2Cre mice, respectively, the DA was released in L5 and L2/3 in response to joystick movement and reward delivery (**Figures 3A and 3B**). As the animals progressed through training, the DA response to reward in L5 diminished, and a significant difference between movement- and reward-related amplitudes became apparent (**Figure 3A**). The D1+ and D2+ populations were likewise simultaneously recruited during joystick movement, and the reward-related component was also present in the Ca^2+^ signals (**Figures 3C and 3D**), suggesting that at the population level, D1+ and D2+ cells in the M1 encode performance outcome. Over time, the reward-related Ca^2+^ response was almost eliminated, and since the motor-related activity was more prominent in the Ca^2+^ data, this would indicate that the D1+ and D2+ populations are primarily involved in the forelimb movement generation. The behavioral data (**Figures 3E and 3F**) revealed that mice licked more vigorously (with higher frequency) at the time of reward delivery (1 to 2 s), but as seen in the ‘late’ sessions, animals from both groups showed the emergence of anticipatory licking, that preceded reinforcement delivery (0 to 1 s). This shows that their behavior was goal-directed and oriented toward attaining a reward. Taken together, this suggest that variations in the reward-related component of DA and Ca^2+^ signals could reflect a change in reward expectation during skill acquisition. Overall, we found that the generation of skilled forelimb movements and reward delivery are encoded by the DA dynamics and population activity of DA-recipient neurons in L2/3 and L5. As reward expectation increased, so did the vigor of joystick movements and anticipatory licking, whereas the neural response to the reward itself gradually declined.

**Figure 3.**
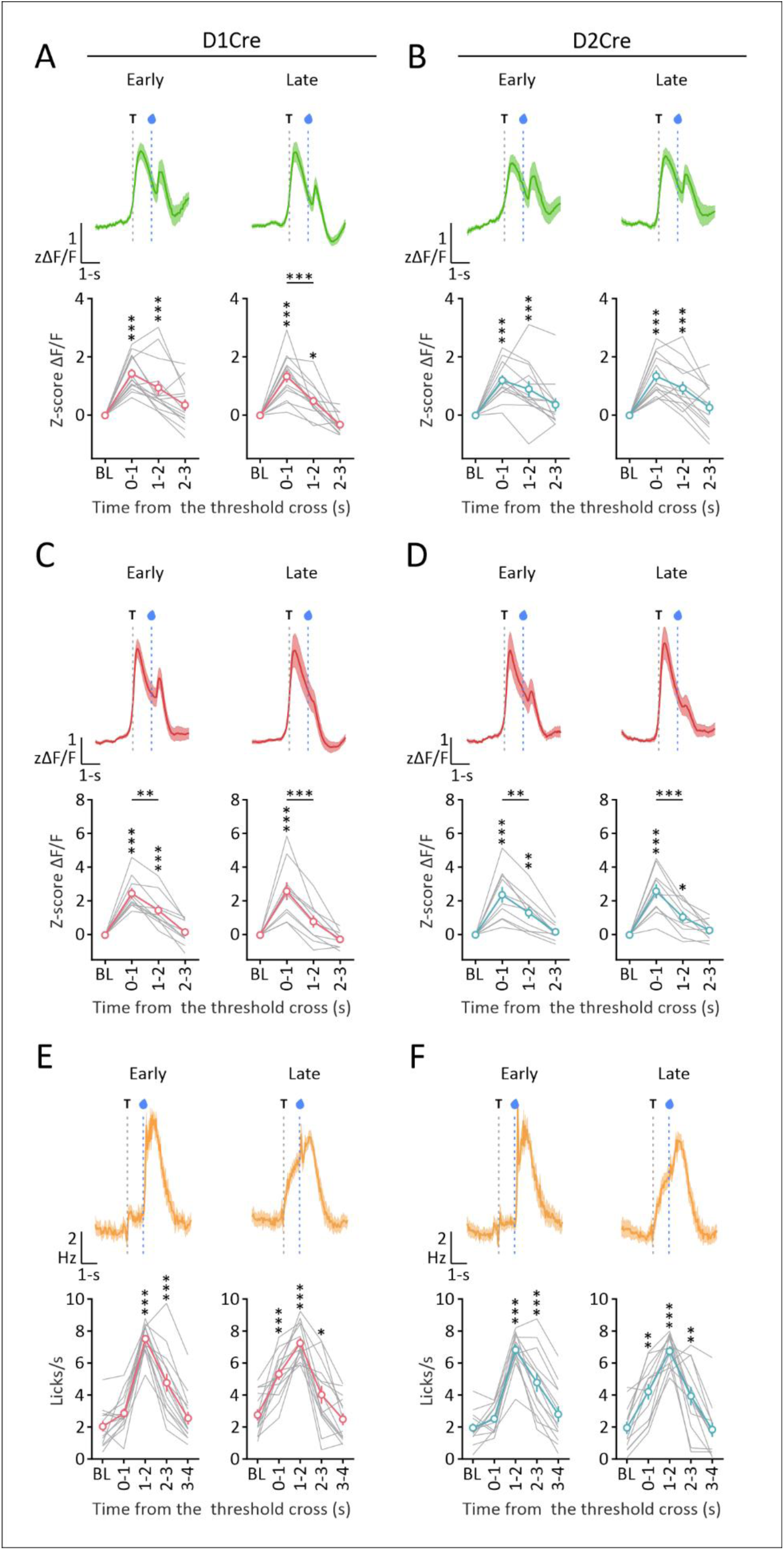
DA dynamics in the M1 encode execution of skilled forelimb movement and reward consumption. (A) Z scored GRAB_DA2h_ fluorescence from L5 in D1Cre mice (n = 14). (B) Z scored GRAB_DA2h_ fluorescence from L2/3 in D2Cre mice (n = 13). (C) Z scored jRGECO1a fluorescence from D1+ neurons in D1Cre mice (n = 10). (D) Z scored jRGECO1a fluorescence from D2+ neurons in D2Cre mice (n = 10). (E) Licking frequency in D1Cre mice (n = 14). (F) Licking frequency in D2Cre mice (n = 13). (A-D) Peri-event plots of fluorescence signals aligned to the threshold crossing (top) and fluorescence signals averaged in time-bins (bottom). (E and F) Peri-event plots of licking behavior aligned to the threshold crossing (top) and lick rates averaged in time-bins (bottom). (A-F) Data are represented as mean ± SEM. Repeated-measure one-way ANOVA; multiple comparisons: baseline (BL) mean vs. every other mean, ‘movement’ (0-1 s) mean vs ‘reward’ (1-2 s) mean; Bonferroni’s post hoc test; \*\*\**p* < 0.001, \*\**p* < 0.01, \**p* < 0.05. Dashed lines indicate joystick movement (threshold crossing) and reward delivery.

### Animals adjust their reach amplitude and the neural response in the M1 scales with the movement vigor

To further test if DA-dependent signaling in the M1 correlates with movement vigor, we subjected the expert mice to additional training session in which the amplitude threshold required to obtain a reward was increased by 3 mm. Although mice in both groups were able to adjust their reach amplitude to the changing threshold requirements, there was no change in movement velocity (**Figures 4A and 4B**). As the increased threshold demanded more effort, we observed a decline in the number of rewarded trials. When we compared the mean fluorescence recorded during joystick movement (0 to 1 s) and reward delivery (1 to 2 s) there were no discernible variations in the DA release between sessions with regular and increased thresholds (**Figures 4C and 4D**). While the D1+ population’s movement-related activity somewhat decreased in the elevated threshold session, the response of the D2+ population remained unchanged between these two conditions (**Figures 4E and 4F**).

**Figure 4.**
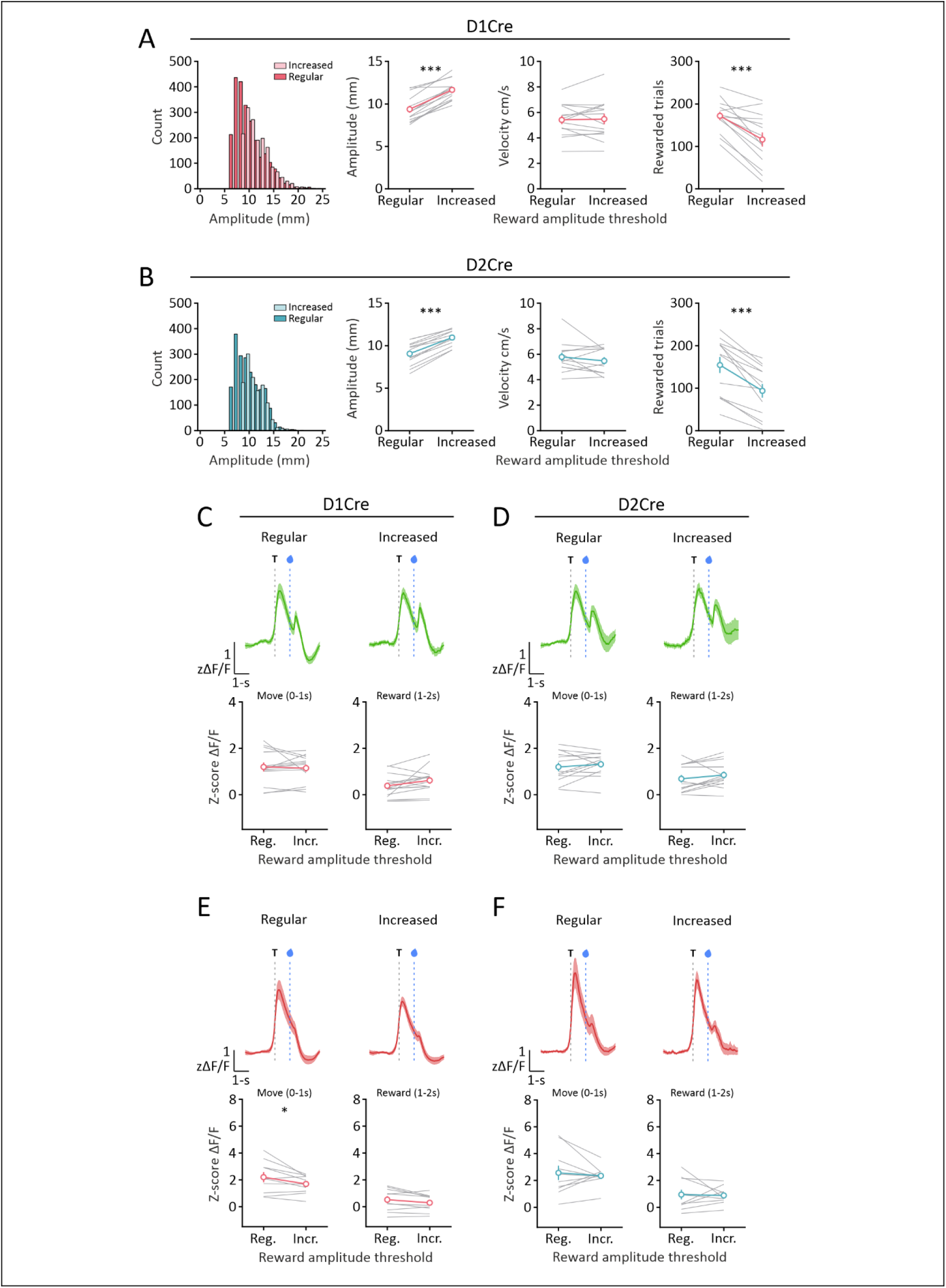
Animals adjust their reach amplitude to the changing threshold requirements. (A) Performance of D1Cre mice in ‘regular’ and ‘increased’ threshold sessions. (Left to right) Distribution of movement amplitudes; movement amplitude; joystick velocity at the time of threshold crossing; number of rewarded movements (n = 14). (B) Performance of D2Cre mice in ‘regular’ and ‘increased’ threshold sessions. (Left to right) Distribution of movement amplitudes; movement amplitude; joystick velocity at the time of threshold crossing; the number of rewarded movements (n = 13). (C) Z scored GRAB_DA2h_ fluorescence from L5 in D1Cre mice (n = 14). (D) Z scored GRAB_DA2h_ fluorescence from L2/3 in D2Cre mice (n = 13). (E) Z scored jRGECO1a fluorescence from D1+ neurons in D1Cre mice (n = 10). (F) Z scored jRGECO1a fluorescence from D2+ neurons in D2Cre mice (n = 10). (C-F) Peri-event plots of fluorescence signals aligned to the threshold crossing (top) and fluorescence signals averaged in time-bins (bottom). (A-F) Data are represented as mean ± SEM. Paired t-test; \*\*\**p* < 0.001, \*\**p* < 0.01, \**p* < 0.05.

Since the number of rewarded attempts was reduced and animals did not produce higher velocity movements, we considered the results of the increased threshold experiment insufficient to draw conclusions regarding the movement vigor. In order to investigate this further, we separated the individual rewarded joystick movements produced during the regular threshold session into low (25% percentile) and high (75% percentile) amplitude movements. In this case, high-amplitude movements were also the movements made at higher velocities (**Figures 5A and 5B**). We found that high-amplitude/high-velocity movements boosted DA levels in the L5 and L2/3 (**Figures 5C and 5D**). These more vigorous movements were also linked to a greater recruitment of D1+ and D2+ populations (**Figures 5E and 5F**). Overall, we discovered that DA dynamics and population activity of DA-receptive neurons in the M1 scaled with the vigor of forelimb movements. Additionally, we found that movement velocity combined with its amplitude was a more reliable predictor of movement vigor and that altering task demands by simply increasing the amplitude threshold was insufficient to induce more vigorous movements.

**Figure 5.**
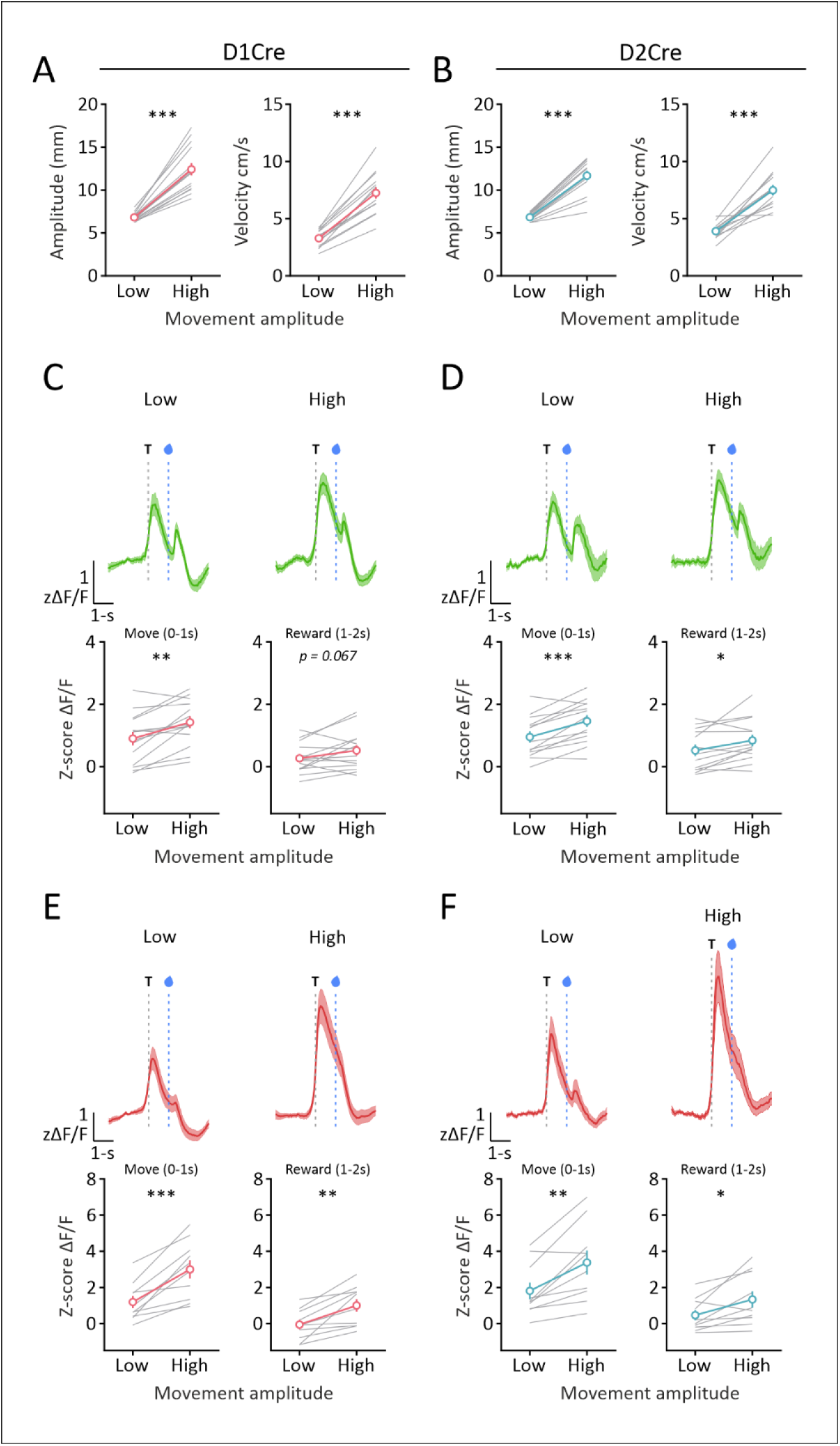
The neural response in the M1 scales with the movement vigor. (A) Performance of D1Cre mice in ‘low’ and ‘high’ amplitude trials. (Left to right) Movement amplitude; joystick velocity at the time of threshold crossing (n = 14). (B) Performance of D2Cre mice in ‘low’ and ‘high’ amplitude trials. (Left to right) Movement amplitude; joystick velocity at the time of threshold crossing (n = 13). (C) Z scored GRAB_DA2h_ fluorescence from L5 in D1Cre mice (n = 14). (D) Z scored GRAB_DA2h_ fluorescence from L2/3 in D2Cre mice (n = 13). (E) Z scored jRGECO1a fluorescence from D1+ neurons in D1Cre mice (n = 10). (F) Z scored jRGECO1a fluorescence from D2+ neurons in D2Cre mice (n = 10). (C-F) Peri-event plots of fluorescence signals aligned to the threshold crossing (top) and fluorescence signals averaged in time-bins (bottom). (A-F) Data are represented as mean ± SEM. Paired t-test; \*\*\**p* < 0.001, \*\**p* < 0.01, \**p* < 0.05.

### The DA in the M1 is sensitive to changes in action-reward contingencies

We were able to temporally isolate the DA-release and neural activity related to reward collection from the modulation of activity during forelimb movement. To further determine whether the neural activity in the M1 is sensitive to changes in action-outcome contingencies, we modified the task to introduce reward delay and omission. First, we trained mice in a session in which the reward was delayed by 3 s. We found that the DA response in L5 and L2/3 decayed at the time of the expected reward (1-2 s) and ramped up again following the actual reward (3-4 s) (**Figures 6A and 6B**). Although primarily recruited during joystick movement, the D1+ and D2+ populations also showed a residual response to delayed reward, suggesting they were sensitive to unexpected shifts in the action-outcome contingency (**Figures 6C and 6D**). Mice from both groups increased their anticipatory licking during the expected reward period (1-2 s) and then further elevated their licking frequency at the time of reward delivery (3-4 s) (**Figures 6E and 6F**).

**Figure 6.**
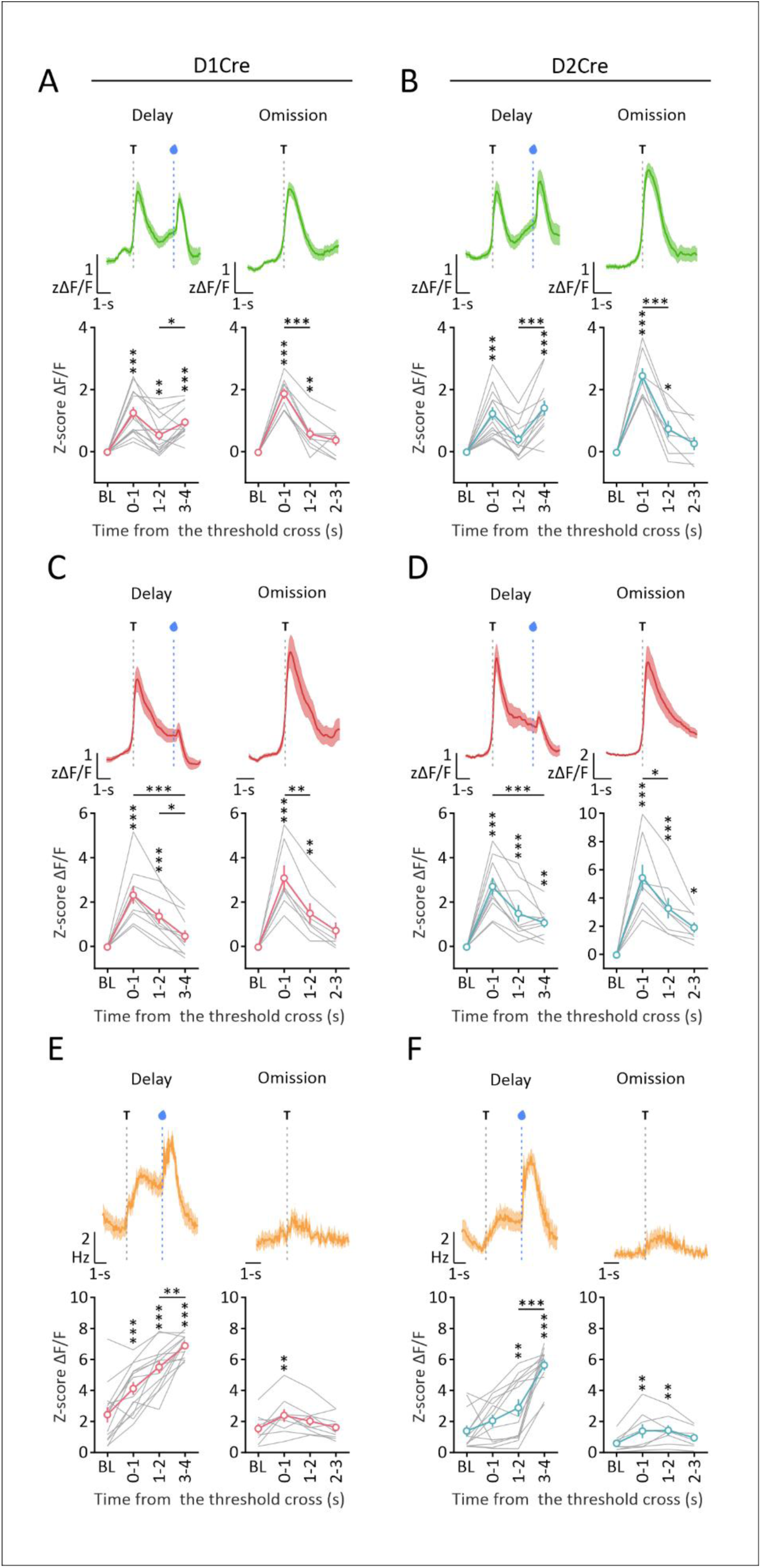
The DA in the M1 is sensitive to changes in action-reward contingencies. (A) Z scored GRAB_DA2h_ fluorescence from L5 in D1Cre mice during ‘delay’ (n = 14) and ‘omission’ (n = 9). (B) Z scored GRAB_DA2h_ fluorescence from L2/3 in D2Cre mice during ‘delay’ (n = 13) and ‘omission’ (n = 8). (C) Z scored jRGECO1a fluorescence from D1+ neurons in D1Cre mice during ‘delay’ (n = 10) and ‘omission’ (n = 7). (D) Z scored jRGECO1a fluorescence from D2+ neurons in D2Cre mice during ‘delay’ (n = 10) and ‘omission’ (n = 8). (E) Licking frequency in D1Cre mice during ‘delay’ (n = 14) and ‘omission’ (n = 9). (F) Licking frequency in D2Cre mice during ‘delay’ (n = 13) and ‘omission’ (n = 8). (A-D) Peri-event plots of fluorescence signals aligned to the threshold crossing (top) and fluorescence signals averaged in time-bins (bottom). (E and F) Peri-event plots of licking behavior aligned to the threshold crossing (top) and lick rates averaged in time-bins (bottom). (A-F) Data are represented as mean ± SEM. Repeated-measure one-way ANOVA; multiple comparisons: baseline (BL) mean vs. every other mean; ‘movement’ (0-1 s) mean vs ‘actual-reward’ (3-4 s) mean; ‘expected-reward’ (1-2 s) mean vs ‘actual-reward’ (3-4 s) mean Bonferroni’s post hoc test; \*\*\**p* < 0.001, \*\**p* < 0.01, \**p* < 0.05. Dashed lines indicate joystick movement (threshold crossing) and reward delivery.

Secondly, a subset of mice was trained under a schedule in which the expected reward was completely omitted. We found that the reward-related component of the signal in both DA (**Figures 6A and 6B**) and Ca2+ (**Figures 6C and 6D**) data was absent, and DA levels and population activity of D1+ and D2+ cells elevated by the joystick movement slowly returned to baseline. Although mice were continuously licking, their lick rate oscillated at a low rate (∼2 Hz) near the baseline level (**Figures 6E and 6F**). Overall, we showed that DA release and population activity of DA-recipient neurons in the M1 tracked the relationship between actions and their consequences. We also confirmed that animals’ behavior remained goal-directed, even after more than 3 weeks of training, as they adapted their performance to the degradation of the action-reward contingency.

### The release of photoinhibition of DA-sensitive neuronal populations in the M1 reduces the likelihood of rewarded joystick movement

Next, we asked whether DA-sensitive populations in the M1 are necessary for the execution of learned forelimb movements in the joystick task. We induced expression of archaerhodopsin T (ArchT)^28^, a light-driven outward proton pump that causes membrane hyperpolarization, in D1+ and D2+ populations in the M1 (**Figures 7A and 7E**). No ArchT-expressing (tdTomato+) cells were detected in control (Cre-) mice (**Figure 7I**). Photoinhibition experiments were carried out when animals had completed 12 training sessions (**Figures 7B, 7F and 7J**) and involved bilateral intracranial light delivery (561 nm) for 5 min, preceded and followed by 5 min of laser-Off time (**Figures 7C, 7G and 7K**). The light intensity was increased between sessions from 1.5 mW to 6 mW and 10 mW. We also examined mice performance during a session in which the laser light was completely turned Off, and a session in which the reward threshold was increased by 3 mm. Although we did not observe the direct effects of photoinhibition during the laser-On period, we found that the subsequent laser-Off period was associated with a reduction in rewarded trials in D1Cre and D2Cre animals, and increasing laser intensity appeared to amplify the effect (**Figures 7C and 7G)**. However, a similar, although smaller, response was observed during the laser-Off session, and in the control group (**Figure 7K**). But the response of D1Cre animals to the increased threshold, when they gathered fewer rewards in the previous time blocks, may confirm that the effect was attributed to the release of inhibition and not simply satiation or a decrease in motivation. There was no consistent effect of photoinhibition on movement amplitude or velocity (**Figures 7C, 7G and 7K**). Additionally, we did not find significant variations in movement trajectories between blocks with and without light (**Figures 7D, 7H and 7L**). Overall, we found that the number of rewarded trials decreased following photoinhibition of the D1+ and D2+ populations in the M1 (during the last five minutes of the session), but there was no discernible change in movement amplitude, velocity, or direction.

**Figure 7.**
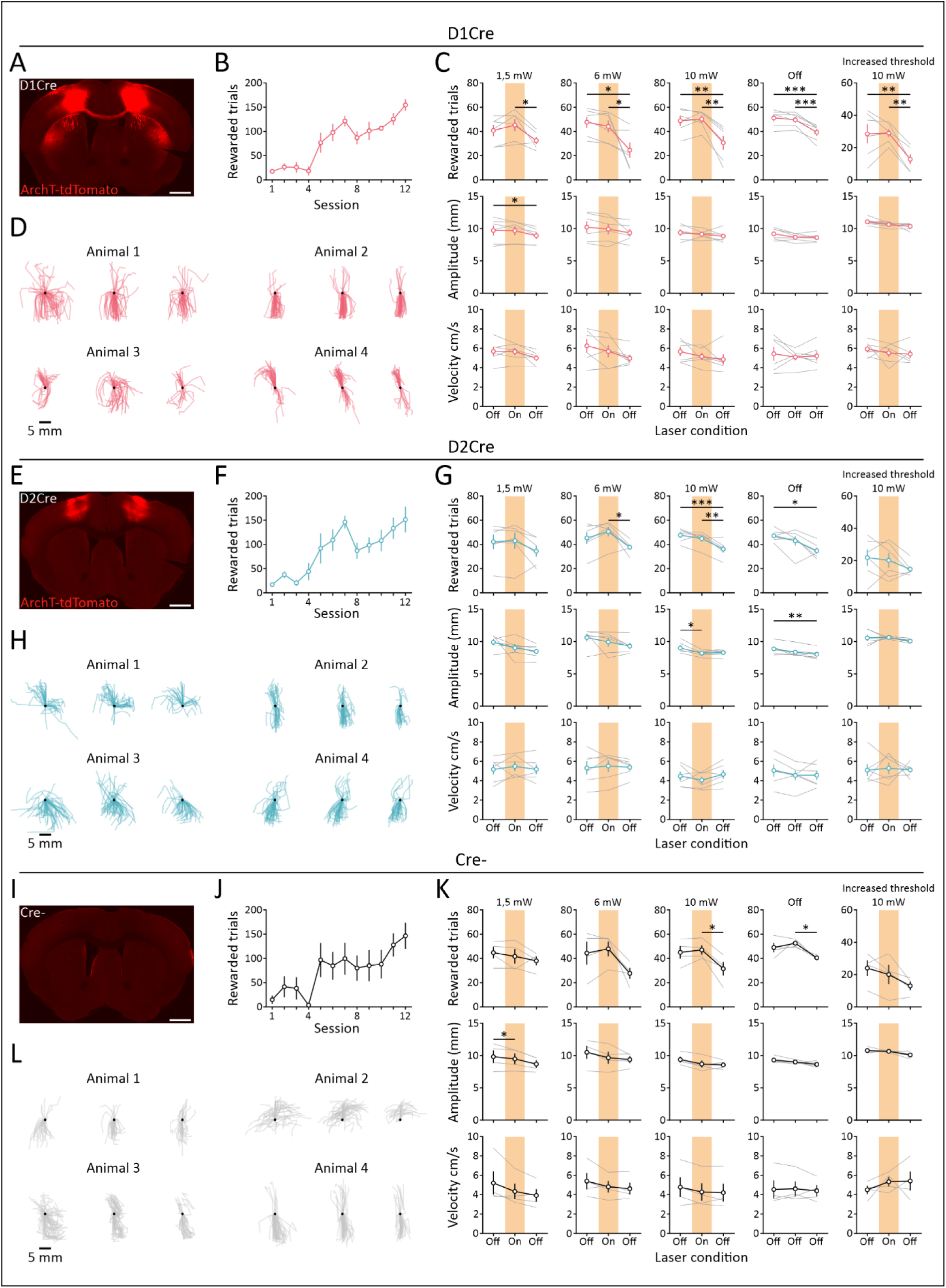
The release of photoinhibition of DA-sensitive neuronal populations in the M1 reduces the likelihood of rewarded joystick movement. (A) Coronal section obtained from a D1Cre mouse showing expression of ArchT-tdTomato. Scale bar, 500 µm. (B) The number of rewarded joystick movements during training in D1Cre mice (n = 6). (C) Performance of D1Cre mice in photoinhibition sessions. (Top to bottom) The number of rewarded movements; movement amplitude; joystick velocity at the time of threshold crossing (n = 6). (D) Movement trajectories of representative D1Cre mice aligned to the initial joystick position (black dot). (E) Coronal section obtained from a D2Cre mouse showing expression of ArchT-tdTomato. Scale bar, 500 µm. (F) The number of rewarded joystick movements during training in D2Cre mice (n = 6). (G) Performance of D2Cre mice in photoinhibition sessions. (Top to bottom) number of rewarded movements; movement amplitude; joystick velocity at the time of threshold crossing (n = 6). (H) Movement trajectories of representative D2Cre mice aligned to the initial joystick position (black dot). (I) Coronal section obtained from a Cre-mouse showing a lack of ArchT-tdTomato expression. Scale bar, 500 µm. (J) The number of rewarded joystick movements during training in Cre-mice (n = 4). (K) Performance of Cre-mice in photoinhibition sessions. (Top to bottom) number of rewarded movements; movement amplitude; joystick velocity at the time of threshold crossing (n = 4). (L) Movement trajectories of representative Cre-mice aligned to the initial joystick position (black dot). (B and C, F and G, J and K) Data are represented as mean ± SEM. (C, G and K) Paired t-test; \*\*\**p* < 0.001, \*\**p* < 0.01, \**p* < 0.05.

### A subset of D1+ and D2+ neurons of the M1 contacts long-range targets

We previously traced the axonal projections of D1+ and D2+ cells in the M1 using an anterograde tracing approach and found labeled axons in various cortical and subcortical brain regions^19^. Here, we wanted to confirm specific long-range connections of D1+ and D2+ neurons. We used D1-tdTomato^29^ (**Figure 8**) and D2Cre::Ai14 (td-tomato)^26,30^ (**Figure 9**) mice and injected rAAV2-retro-hSyn-EYFP (rAAV2-retro)^31^ or Fluoro-Green (FGr)^32^ retrograde tracers (which labeled cells in green) at specific target locations including the contralateral M1 (cM1), ipsilateral and contralateral dorsal striatum (iDLS and cDLS), ipsilateral thalamus (iThal), and ipsilateral pontine nucleus (iPN) (**Figures 8A and 9A**). We obtained slices containing the M1 forelimb area (A/P -0.10 to 1.00 from the bregma) and counted D1+ and D2+ (tdTomato-positive), EYFP+ (EYFP-positive), and FGr+ (FGr-positive) cells. We plotted the laminar distribution of D1+ and D2+ cells together with the projection-specific distributions of retrogradely labeled EYFP+ and FGr+ neurons (**Figures 8B and 9B**). Consistent with our previous observation^19^, D1+ neurons were most heavily concentrated in deep layers (L5-6), with lower relative abundance in L2/3 (**Figures 8B and S1**). In contrast, D2+ cells were concentrated in the upper L2/3, with relatively low expression below (**Figures 9B and S1**). Next, we examined the colocalization of D1+ or D2+ neurons with different populations of EYFP+ and FGr+ projection neurons. We quantified the percentage of D1+ neurons that are also EYFP+ (**Figure 8C**, % (D1+ EYFP+)/D1+), and D1+ neurons that are also FGr+ (**Figure 8D**, % (D1+ FGr+)/D1+), and did the same with D2+ neurons (**Figures 9C and 9D**). In each case, we found relatively strong colabeling for cells projecting to cM1 (∼20 % in D1+ and ∼10

**Figure 8.**
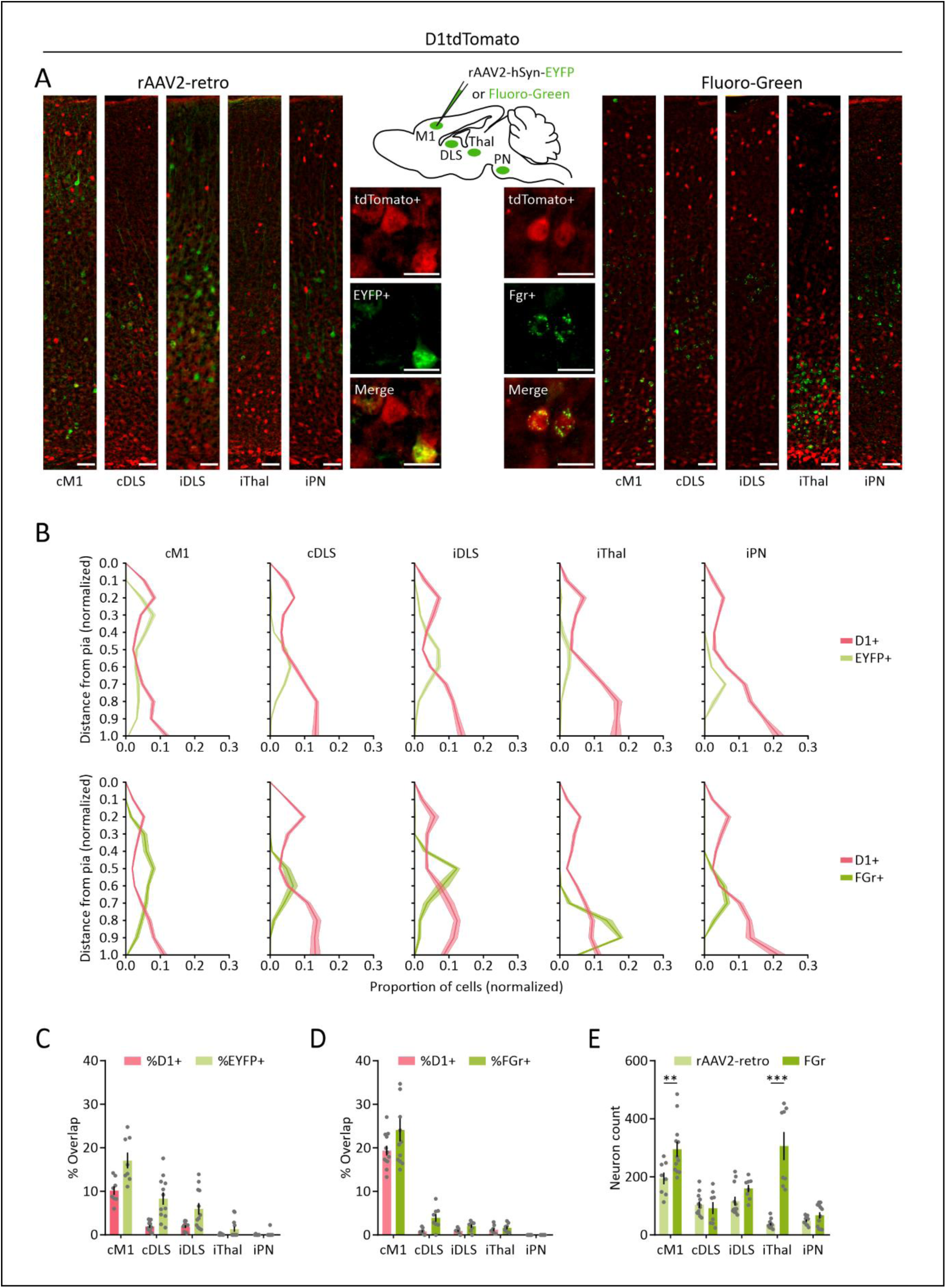
A subset of D1+ neurons of the M1 contacts long-range targets. (A) Schematic of the retrograde tracers injection sites and representative coronal sections obtained from D1tdTomato mice showing laminar distribution of tdTomato expressing D1+ neurons and EYFP or FGr expressing retrogradely labeled projection neurons. Insets show representative colocalization between tdTomato expressing D1+ neurons and retrogradely labeled cells. Scale bars, main image 50 µm, inset 20 µm. (B) Normalized cell density of D1+ and projection neurons as a function of distance from pia (0, pia; 1, white matter); cell number per unit normalized to a total number of fluorescently labeled cells within the section. (C) Percentage of D1+ neurons and the percentage of EYFP+ neurons. (D) Percentage of D1+ neurons and the percentage of FGr+ neurons. (E) Comparison of labeling efficacy of rAAV2-retro and FGr tracers expressed by the number of labeled cells. (B-E) Data are represented as mean ± SEM. rAAV2-retro: cM1 (n = 2), cDLS (n = 3), iDLS (n = 3), iThal (n = 3), iPN (n = 3); n = 4 slices per animal. FGr: cM1 (n = 3), cDLS (n = 2), iDLS (n = 2), iThal (n = 2), iPN (n = 3); n = 4 slices per animal. (E) Two-way ANOVA; multiple comparisons: EYFP mean vs FGr mean; Bonferroni’s post hoc test; \*\*\**p* < 0.001, \*\**p* < 0.01. See also **Figure S1**.

**Figure 9.**
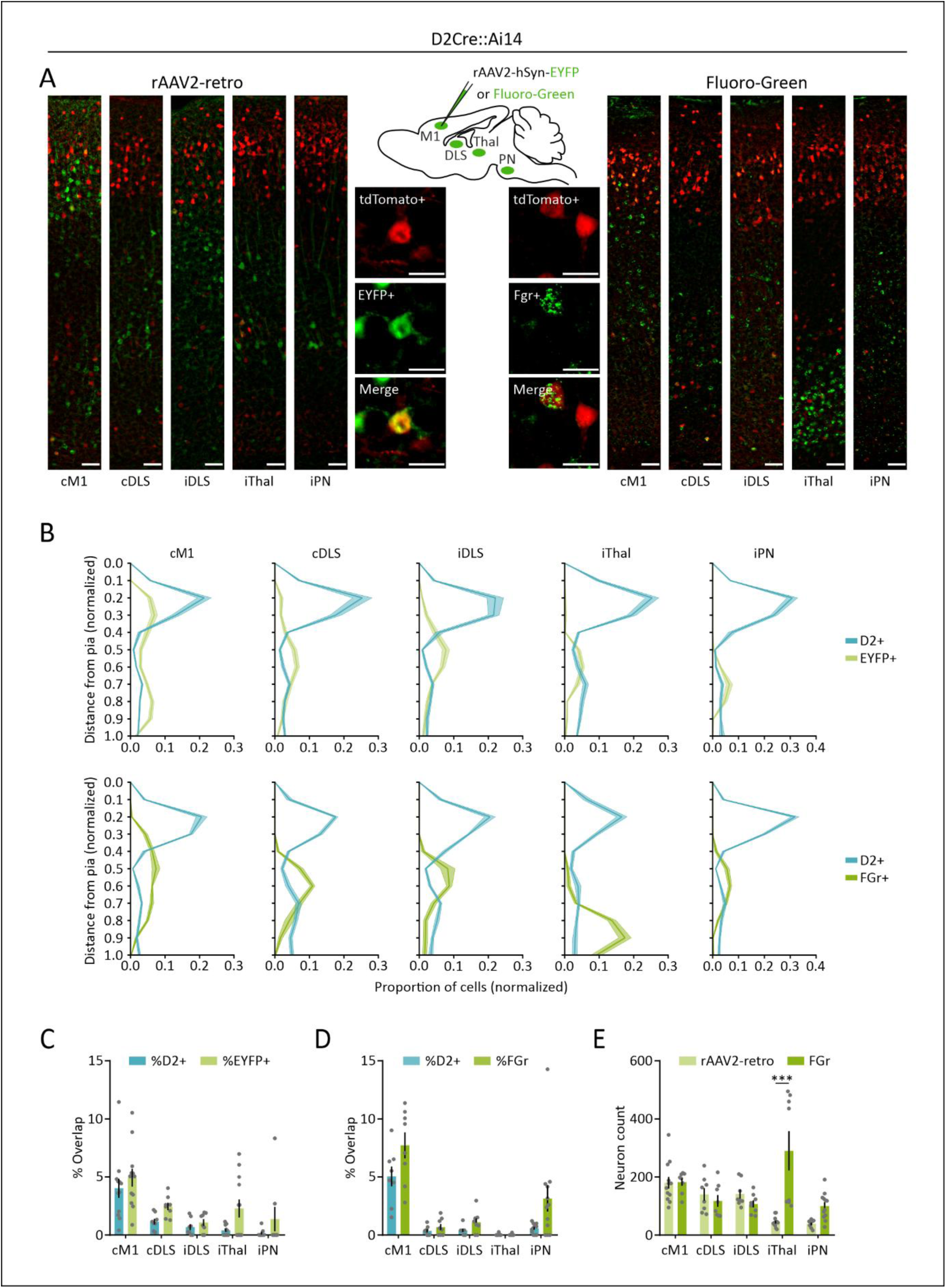
A subset of D2+ neurons of the M1 contacts long-range targets. (A) Schematic of the retrograde tracers injection sites and representative coronal sections obtained from a D2Cre::Ai14 mice showing laminar distribution of tdTomato expressing D2+ neurons and EYFP or FGr expressing retrogradely labeled projection neurons. Insets show representative colocalization between tdTomato expressing D2+ neurons and retrogradely labeled cells. Scale bars, main image 50 µm, inset 20 µm. (B) Normalized cell density of D2+ and projection neurons as a function of distance from pia (0, pia; 1, white matter); cell number per unit normalized to a total number of fluorescently labeled cells within the section. (C) Percentage of D2+ neurons and the percentage of EYFP+ neurons. (D) Percentage of D2+ neurons and the percentage of FGr+ neurons. (E) Comparison of labeling efficacy of rAAV2-retro and FGr tracers expressed by the number of labeled cells. (B-E) Data are represented as mean ± SEM. rAAV2-retro: cM1 (n = 3), cDLS (n = 2), iDLS (n = 2), iThal (n = 3), iPN (n = 2); n = 4 slices per animal. FGr: cM1 (n = 2), cDLS (n = 2), iDLS (n = 2), iThal (n = 2), iPN (n = 3); n = 4 slices per animal. (F) Two-way ANOVA; multiple comparisons: EYFP mean vs FGr mean; Bonferroni’s post hoc test; \*\*\**p* < 0.001. See also **Figure S1**.

% in D2+ population), with much less colabeling for cells projecting to other regions (less than 5 %). Because the ratios of tdTomato+ to retrogradely labeled cells were not equal, this analysis might have underestimated the number of colabeled cells. Therefore, we also quantified the percentage of EYFP+ neurons that are D1+ (**Figure 8C**, % (D1+ EYFP+)/EYFP+), and FGr+ neurons that are D1+ (**Figure 8D**, % (D1+ FGr+)/FGr+), and did the same with D2+ neurons (**Figures 9C and 9D**). In this case, the overlap was slightly higher, but the percentage of colabeled neurons projecting to subcortical targets was still relatively low, showing that only a small fraction of neurons from D1+ and D2+ populations contact long-range targets.

Although the laminar distributions of retrogradely labeled cells were comparable between two tracers, we observed a difference in the laminar distribution of iThal projecting cells (**Figures 8B and 9B**). We speculated that the rAAV2-retro and FGr had different labeling efficacies, which could account for the observed effect. To further test this, we compared the total number of neurons labelled with the use of both methods. The analysis revealed that the FGr-injected mice from the D1tdTomato and D2Cre::Ai14 strains did have a greater number of iThal labeled neurons than the rAAV2-retro-injected animals (**Figures 8E and 9E**). A considerably greater amount of cM1 labeled neurons in FGr-injected animals was also confirmed in D1tdTomato strain (**Figure 8E**). Overall, we confirmed that D1+ and D2+ neurons have discreate distribution in the layers of the motor cortex. We also showed that only a small fraction of D1+ and D2+ projection neurons of the M1 contact long-range targets. However, the reported effects may have been affected by the labeling efficacies of the retrograde tracers used in the study.

## Discussion

Previous studies focusing on the long-term consequences of DA depletion have shown that dopaminergic input to M1 forelimb area is necessary for the acquisition of motor skills and modulation of M1 plasticity^6,11,16^. Here, we monitored the DA release across the M1 layers 2/3 and 5, and recorded the activity of D1+ and D2+ populations in D1Cre and D2Cre mice trained in a forelimb-specific joystick task. We found that DA release events and Ca^2+^ transients were temporally linked to joystick movements (threshold crossing) and reward consumption. The DA signal was uniformly broadcast across the superficial and deep layers of M1. Likewise, both populations of D1+ and D2+ neurons were recruited concomitantly during movement onset. As training progressed, the difference in DA signal amplitudes associated with movement and reward became noticeable, particularly in L5, likely indicating a change in reward expectation as seen in anticipatory licking. Moreover, we found that the DA response tracked the delay in reward delivery and was absent following reward omission. Interestingly, dopaminoceptive populations of M1 were also responsive to rewards, although the magnitude of reward-related Ca^2+^ response was much smaller than in the case of DA. This is consistent with the general role of DA in reinforcement learning^33,34^, and a recent observation showing that the M1 neurons represent the outcome of movement performance, such as success or failure, and that DA input to M1 is crucial for outcome encoding and skill acquisition^35,36^. Together with our findings, it would indicate that in addition to the previously reported role of DA in modulating long-term plasticity, the phasic DA signaling in M1 also acts as a temporary teaching signal supporting reinforcement motor learning of skilled behavior.

The ability to execute motions over a range of amplitudes and speeds, known as movement vigor, is another aspect of motor skill learning, and it is generally accepted that DA invigorates and motivates movement^37,38^. In line with this, we found that high-amplitude/high-velocity movements resulted in elevated DA release and greater recruitment of both dopaminoceptive populations. However, when we raised the difficulty of the task by increasing the amplitude threshold required to trigger the reward, we found that forelimb movement velocities remained unchanged and task engagement, measured by a number of rewarded trials, declined. Furthermore, neither the amplitude nor the speed of forelimb movement was significantly impacted by photoinhibition of the D1+ or D2+ populations. This would suggest that while the decision to generate vigorous movement is associated with the elevation of DA level in M1, the gain of motor commands is regulated elsewhere, most likely in the basal ganglia^37,39,40^.

As shown previously, motor cortex is required for learning but not execution of motor skills^41^. However, Guo et al.’s^1^ seminal research showed that abrupt cortical inhibition prevents the initiation and execution of a learned skilled prehension behavior, proving that M1 activity is necessary for performing learned skilled forelimb movements. Therefore, we employed reversible optogenetic inhibition to probe the role of DA-sensitive motor cortical circuits in our joystick task. Although photoinhibition had no direct influence on motor performance, we found that when cortical inhibition was released, the number of rewarded joystick movements in D1Cre and D2Cre animals decreased. The reduced probability of movement initiation reflected the effects of cortical inhibition in the M1 forelimb area, which was accomplished by injecting GABAA agonist (muscimol)^42,43^, or stimulating cortical inhibitory interneurons expressing the vesicular GABA transporter (VGAT)^1,21,44^. In the neocortex, D1 and D2 receptors are expressed in both pyramidal and GABAergic interneurons^17,18,45^ and as shown in our previous research^19^, both populations of D1+ and D2+ cells contain ∼20% of VGAT+ GABAergic cells. Therefore, we hypothesize that the effects shown in the photoinhibition experiment result from GABAergic interneuron rebound activity. Given that excessive ArchT stimulation can result in rebound hyperactivation of ArchT-expressing cells^46^ and that Guo et al. observed prominent rebound spiking in cortical neurons after prolonged photostimulation, this explanation seems plausible. However, the precise mechanism is difficult to determine due to the diversity of D1+ and D2+ populations, necessitating further investigation.

Axonal projections from the mouse motor cortex have been extensively studied, revealing a very complex network^3,47–50^. We have previously provided details of the laminar distribution of D1+ and D2+ somas in the M1 forelimb area and traced their axonal projections^19^. Here, we injected retrograde tracers into previously identified regions of axonal projections, to determine the long-range connections of D1+ and D2+ projection neurons in more detail. Among the identified long-range connections, the majority were made with the contralateral cortex, and to a lesser extent with the striatum, thalamus and pons. Nevertheless, even though both tracers were effective in retrograde labeling of projection neurons, we found a relatively low degree of colocalization between retrogradely labeled cells and tdTomato expressing D1+ and D2+ neurons. We also found that the FGr tracer was more effective in labeling cortico-thalamic projections than AAV2-retro. This would suggest that either a small percentage of D1+ and D2+ cells are in fact projection neurons or that their number was underestimated. Despite the fact that the AAV2-retro was demonstrated to be an effective retrograde viral tracer, it was suggested that the labeling efficacy of cortico-thalamic projections is relatively weak, while that of cortico-striatal projections is moderate^31^. Furthermore, previous studies utilized rabies or classic retrograde tracers (fluorogold and cholera toxin B subunit) to define the projections from the M1 forelimb area^3,49,50^. Therefore, we can’t exclude the possibility that the use of different tracers would provide better results.

Overall, we have demonstrated for the first time that the DA dynamics and activity of local dopaminoceptive circuits in M1 encode voluntary forelimb movement and it’s vigor, reward, and change in the action-outcome contingency. Our findings fill a gap in understanding how phasic DA signals in M1 facilitate reinforcement motor learning of skilled behavior.

## Resource availability

### Lead contact

Any additional information required to reanalyze the data reported in this paper is available from the lead contact upon request.

### Materials availability

This study did not generate new unique reagents. Headbar design for head-fixation will be made available upon request. Joystick built protocol and Arduino source code was provided by Yttri Lab (https://github.com/YttriLab/Joystick).

### Data and code availability

Python code for data analysis reported in this paper is provided by Gniewosz Drwiega (https://github.com/gniewko-d/Joystick_task/)

## Acknowledgments

We would like to thank Alex Kwan (Department of Psychiatry, Yale University School of Medicine, New Haven, CT, USA) for sharing the headbar design for head-fixation. This research was funded in whole by the National Science Centre, Poland – grant SONATA 2020/39/D/NZ4/00503 (to P.E.C.). G.D. was supported by the grant PRELUDIUM 2022/45/N/NZ4/03171.

## Author contributions

P.E.C. designed the study and wrote the manuscript. M.G-N. and P.E.C. performed the experiments and analyzed data. G.D. wrote code for data analysis. J.R.P. provided transgenic mice. L.S. genotyped animals and managed the colony. M.G.-N., G.D. and J.R.P. revised and edited manuscript.

## STAR Methods

### Experimental model and study participant details

#### Animals

All animal procedures were approved by the 2nd Local Institutional Animal Care and Use Committee in Krakow (approval number 223/2022, issued on 04 August 2022) and conducted in accordance with the Directive 2010/63/EU of the European Parliament, and of the Council of 22 September 2010 on the protection of animals used for scientific purposes. Mice were housed 2 – 5 per cage in an animal facility room with a controlled temperature (22 ± 2°C) and humidity (40-60% RH), under a 12 h light/dark cycle. Unless otherwise specified, mice had *ad libitum* access to water and laboratory chow (VRF1, Altromin). D1Cre mice^25^ were obtained from the German Cancer Research Center, Heidelberg, and D2Cre^26^ from the University of California, Davis (MMRRC_032108-UCD). The D1-tdTomato line 6^29^ and Ai14 (tdTomato) Cre reporter line^30^ were purchased from The Jackson Laboratory (IMSR_JAX: 016204 and IMSR_JAX: 007914). For the purpose of the project, the D2Cre and Ai14 (tdTomato) strains were crossed to obtain D2Cre::Ai14, double transgenic animals. All mice were congenic with the C57BL/6N background (>8 generations of backcrosses prior to initiation of the study). Genotyping was performed using a standard PCR assay according to previously described protocols and genotyping protocols available in the JAX database. A total of *n* = 72 mice of both sexes, aged 8-12 weeks at surgery were used in the experiments.

### Method details

#### Surgery

Mice were anesthetized with a mixture of ketamine (100 mg/ml, Biowet) and xylazine (20 mg/ml, Biowet) and placed into an automated stereotaxic instrument (model 71000, RWD Life Science Co., Ltd.). During surgery, eyes were protected from drying by application of eye drops (Hylo Gel, Ursapharm), and body temperature was maintained at 37°C by an automatic heating pad. After cutting the skin and exposing the skull surface, a small craniotomy above the region of interest was made using a hand drill. Borosilicate glass pipettes (B114-53-10NP, Sutter Instrument) were pulled with a vertical glass microelectrode puller (model PE-21, Narishige) to obtain ∼30 µm tips, backfilled with paraffin oil (76235-500ML, Sigma-Aldrich), and connected to a glass-capillary Nanoinjector (model NanoW, Neurostar GmbH). The adeno-associated virus (AAV) solution or fluorescent emulsion was front-loaded into the pipette and injected into the brain tissue at a rate of 1 nl/s. The pipette was kept in place for 5 min after injection before being slowly withdrawn.

The following viral vectors and fluorescent tracer were used in the experiments: AAV9-hSyn-GRAB ^23^ and AAV1-Syn-Flex-NES-jRGECO1a^24^ for *in vivo* photometry; AAV5-Flex-ArchT-tdTomato^28^ for optogenetic photoinhibition; rAAV2-Retro-hSyn-EYFP^31^ and Fluoro-Green^32^ for retrograde tracing. Viral vectors were obtained from Addgene (140554-AAV9, 100853-AAV1, 28305-AAV5) or UNC GTC Vector Core (rAAV2-retro); Fluoro-Green (FGr) was manufactured by Tombow Pencil Co., Ltd. AAVs were injected at a volume of 100-200 nl, and FGr at a volume of 20-60 nl per site, according to the following coordinates: M1 forelimb area (M1: A/P 0.5 mm; M/L 1.5 mm from the bregma; D/V -0,9 mm [D1Cre] or -0,4 mm [D2Cre] from pia); dorsolateral striatum (DLS: A/P 0.5 mm; M/L 2.5 mm; D/V -3,3 to -3,0 mm from the bregma); thalamus (THAL: A/P -1,4 mm; M/L 1.5 mm; D/V -3,8 to -3,3 mm from the bregma) or pontine nucleus (PN: A/P -4,2 to -4,0 mm; M/L 0.5 mm; D/V -5,6 to -5,4 mm from the bregma).

Fiber-optic cannulas (optical fiber: Ø200 µm, 0.39 NA; ceramic ferrule: Ø230 µm, 1.25 mm/6,4 mm long; Thorlabs) were made in-house and implanted during the same surgery. In the photometry experiment, the tip of the optical fiber was inserted into the right M1 and positioned 150 to 200 µm above the injection site, whereas in the optogenetic experiment, the tips of optical fibers were positioned bilaterally onto the brain’s pia. Following the placement of the optical fibers, two anchor screws and the custom-made stainless steel or 3D-printed headplate were attached to the exposed surface of the skull. Dental adhesive resin cement (Super-Bond Universal, Sun Medical) was used to secure everything in place. After surgery, animals were given subcutaneous injections of analgesic, anti-inflammatory drug Tolfedine 4% (40 mg/ml, Vetoquinol) and 5% glucose solution.

#### Joystick task

The joystick task was designed based on the open-source joystick manipulandum for mice^20^. The behavioral training setup consisted of a custom-made head-fixation platform made of solid aluminum optical breadboards and optical post assemblies (Thorlabs); a control unit and data acquisition board (Arduino Mega 2560 Rev3, Arduino); an SD card reader module; a water delivery circuit controlled by a mini-solenoid valve (12 V, 0,04 MPa); and peripherals for interacting with the animal: a spring-loaded miniature hall effect joystick (TS1, Ruffy Controls) and lick spout (blunt tip needle) coupled to a capacity touch sensor (MPR121, Adafruit) for lick detection. Mouse behavior was recorded using USB 3.0 camera (model MV-SUA502C-T, MindVision) equipped with 5MP fixed 4 mm focal length C-Mount lens (model VM0420MP5, ZLKC); or USB 3.0 camera (acA1440-220uc, Basler) equipped with 5MP 8 mm fixed focal length C-Mount lens (C125-0818-5M, Basler) placed to the left of the animal. Camera captured 90 frames per second at a resolution of 640 by 480 pixels.

Mice were given about a week to recover from surgery before starting water restriction. During this period, animals received 1 ml of water per day and were gradually acclimated to head restraint for another week. After two weeks, head-fixed mice were trained to make self-initiated (uncued) forelimb movements with a joystick to acquire delayed water reward. When the joystick position surpassed the amplitude threshold by 3 mm, a 3 µl drop of saccharin-flavored (0.01% w/v) water was delivered after a 1 s delay, followed by a 3 s inter-trial interval (ITI) in which no movement would be rewarded. The joystick position at the end of the ITI served as the starting point for the next trial (almost always close to the central default location). Head bar length and joystick distance from the platform were increased by 5 mm after the 3^rd^ session, in order to develop the strength and endurance necessary to complete the task. In the initial days, some mice needed assistance from the experimenter to grab and move the joystick; these trials were not included in the analysis. Once the initial connection between joystick movement and reward was established (usually by session 7), the set reward threshold was raised from 3 mm to 6 mm and maintained as the default threshold for the rest of the experiment. After 14 training sessions, the animals went through extra sessions in which the reward threshold was suddenly increased (to 9 mm), and the reward was either delayed (by 3 s) or not given at all (omission). Regular sessions with a 6 mm threshold and a 1 s reward delay were used to break up these unique sessions. Each animal completed up to 25 sessions.

Each time the suprathreshold joystick movement was detected, a TTL signal was sent from the Arduino-controlled behavioral setup to a fiber photometry recording device. The behavioral events (joystick movements and licks) were recorded and stored on a SD card as a CSV file. A custom-made Python script was used to further analyze the training session data offline. Mice were trained once a day, 6 days per week, and behavioral sessions lasted up to 20 min. At the end of the training session, the amount of water that each animal had consumed was calculated, and if required, mice were given extra water to supply 1 ml each day.

#### Fiber photometry recording

Fiber photometry acquisition was performed with a commercial Tricolor Multichannel Fiber Photometry System (model R820, RWD Life Science) equipped with three LED excitation light sources – 560 nm for the red calcium indicator (jRGECO1a), 470 nm for the green dopamine probe (GRAB_DA2h_) and 410 nm for the reference signal. A multi-branched fiber-optic patch cable (2 m long, Ø200 µm, 0.37 NA, RWD Life Science) with two isolated fibers, permitting signal recording in two animals at the same time, was attached to the implanted fiber-optic cannulas with a zirconia sleeve. LEDs were set to provide ∼30 µW of light intensity at the fiber tip; light intensity was kept constant across sessions. Temporally intermingled excitation pulses of alternating wavelengths were delivered to avoid crosstalk between green and red fluorescence, and emissions were collected by two independent CMOS camera detectors at 20 Hz. To minimize the possibility of fluorescence signal bleaching, *in vivo* recording data were collected every other day within 20 min.

A dedicated fiber photometry system software was used for signal acquisition and initial processing (RWD Life Science). Pre-processing of the recorded signals included: (1) baseline-correction, which used an iterative weighted least square algorithm to correct the trend of decreasing fluorescence; and (2) motion-correction, which used the robust linear fit to fit the 410 nm signal to 470 nm or 560 nm signals. The motion-corrected signal was then obtained by subtracting the fitted 410 nm signal from the signal of interest. The change in fluorescence relative to the baseline fluorescence (ΔF/F) was further calculated as ΔF/F = (F-F1)/F0, where F represents the target fluorescence, F1 is the fitted 410 nm data, and F0 is the median of the raw fluorescence of the whole session; in the case of the peri-event analysis, F0 was the median of the raw fluorescence from the time window -2 to 0 s. Signals were further normalized as z-score = (x-mean)/std, where x = ΔF/F.

A built-in peak statistics algorithm was used to detect individual transients, defined as events with amplitudes greater than two times the median absolute deviation (MAD) above the median of the time window from 100 to 1300 s. These criteria generally selected peaks that aligned with a human observer’s judgment for detecting transients and behavioral outcomes (individual forelimb movements). For the purpose of analysis of event-related activity, the fluorescence signals aligned with threshold crossing (T = 0 s) were collected at 50 ms intervals, within the time window of -2 to 6 s. Events were saved to CSV files. A custom-written Python script was used for further processing, including trial sorting, averaging data across animals or sessions, and generating peri-event plots.

#### Photoinhibition

Prior to the start of photoinhibition experiments, the animals were water deprived and trained for two weeks to master the joystick task (following the procedures described above). A multi-branched fiber-optic patch cable (2 m long, Ø200 µm, 0.37 NA, RWD Life Science) with two isolated fibers, permitting bilateral intracortical light delivery, was attached to the implanted fiber-optic cannulas with a zirconia sleeve. A branching patch cable was attached to an optical rotary joint splitter (RJ 1x2, Thorlabs), providing a 50:50 intensity split of input light, which in turn was connected to 561 nm yellow-green DPSS laser and adjustable power supply (model: YL561T8-100FC + ADR-800A, Shanghai Laser & Optics Century Co., Ltd.). Photoinhibition was accomplished using constant delivery of laser light of increasing intensities (1.5 mW, 6 mW and 10 mW), assessed on different sessions. A photoinhibition session began with the laser turned off for the first 5 min, followed by a 5 min photoinhibition period and a 5 min of laser off time. Photoinhibition sessions were interspersed with the sessions where the laser was completely turned off.

#### Histology

Mice were euthanized via intraperitoneal injection of Morbital (Biowet) (sodium pentobarbital 133.3 mg/ml + pentobarbital 26.7 mg/ml) and perfused with ice-cold phosphate-buffered saline (PBS, pH 7.4, Roth) followed by 4% formaldehyde (pH 7.4, POCH). Dissected brains were fixed in 4% formaldehyde for 24h at 4°C. Next, brains were transferred to a 30% sucrose solution for at least 48h at 4°C. Coronal slices (50 or 100 µm) containing the motor cortex and striatum, thalamus, or pons were cut on an automatic vibrating blade microtome (VT1000S, Leica Biosystems) or manual cryostat microtome.

Slices were blocked in PBS-T containing 10% Normal Donkey Serum (017-000-121, Jackson ImmunoResearch) and 0.6% Triton X-100 (Sigma-Aldrich) for 1 h at room temperature; followed by incubation with primary antibody dissolved in PBS-T (2% NDS, 0.3% Triton X-100) in antibody dilution solution 1:1000 overnight at 4°C. The following primary antibodies were used: rabbit anti-RFP (600-401-379, Rockland); rabbit anti-GFP (ab6556, Abcam); chicken anti-GFP (ab13970, Abcam). After primary antibody incubation, the tissue was washed in PBS-T and incubated with secondary antibody in antibody dilution solution 1:400 overnight at 4°C. The following secondary antibodies were used: Cy™3 AffiniPure® F(ab’)2 Fragment Donkey Anti-Rabbit IgG (H+L) (711-166-152, Jackson ImmunoResearch); Alexa Fluor® 488 AffiniPure® F(ab’)2 Fragment Donkey Anti-Rabbit IgG (H+L) (711-546-152, Jackson ImmunoResearch); Alexa Fluor® 488 AffiniPure® F(ab’)2 Fragment Donkey Anti-Chicken IgY (IgG) (H+L) (703-546-155, Jackson ImmunoResearch). The tissue was then washed with PBS and mounted on glass slides using Fluoroshield^TM^ with DAPI (F6057, Sigma-Aldrich). Brain sections obtained from Drd2Cre::Ai14 mice injected with FGr and D1Cre or D2Cre mice injected with ArchT-tdTomato was mounted on glass slides without incubation.

Images were acquired using a Zeiss Axio Imager M2 microscope equipped with an Axiocam 503 mono camera. Whole brain images were acquired with a 5x objective and z-stack images of regions of interest were acquired with a 20x objective. ZEN Microscopy Software (Zeiss) was used for image acquisition and initial preprocessing (stitching and background subtraction). ImageJ (NIH) image analysis software was further used for image processing (cropping, brightness, and color adjustment). Cell counting was performed in a ∼850 µm wide region of interest across the motor cortex, using the Cell Counter plugin for ImageJ.

### Quantification and statistical analysis

Results are presented as mean ± SEM. Statistical analysis was based on the assumption that the samples follow a Gaussian distribution. Student’s t-test was applied for statistical comparisons between the two groups, and ANOVA followed by post-hoc analysis (Bonferroni’s multiple comparison test) was used for analysis with multiple groups and repeated measures were incorporated when appropriate. *P* < 0.05 was considered statistically significant. All statistical analyses were conducted using GraphPad Prism 6 software (GraphPad Software, Inc.). Detailed results of the statistical analysis are presented in **Table 1**.

**Table 1.**
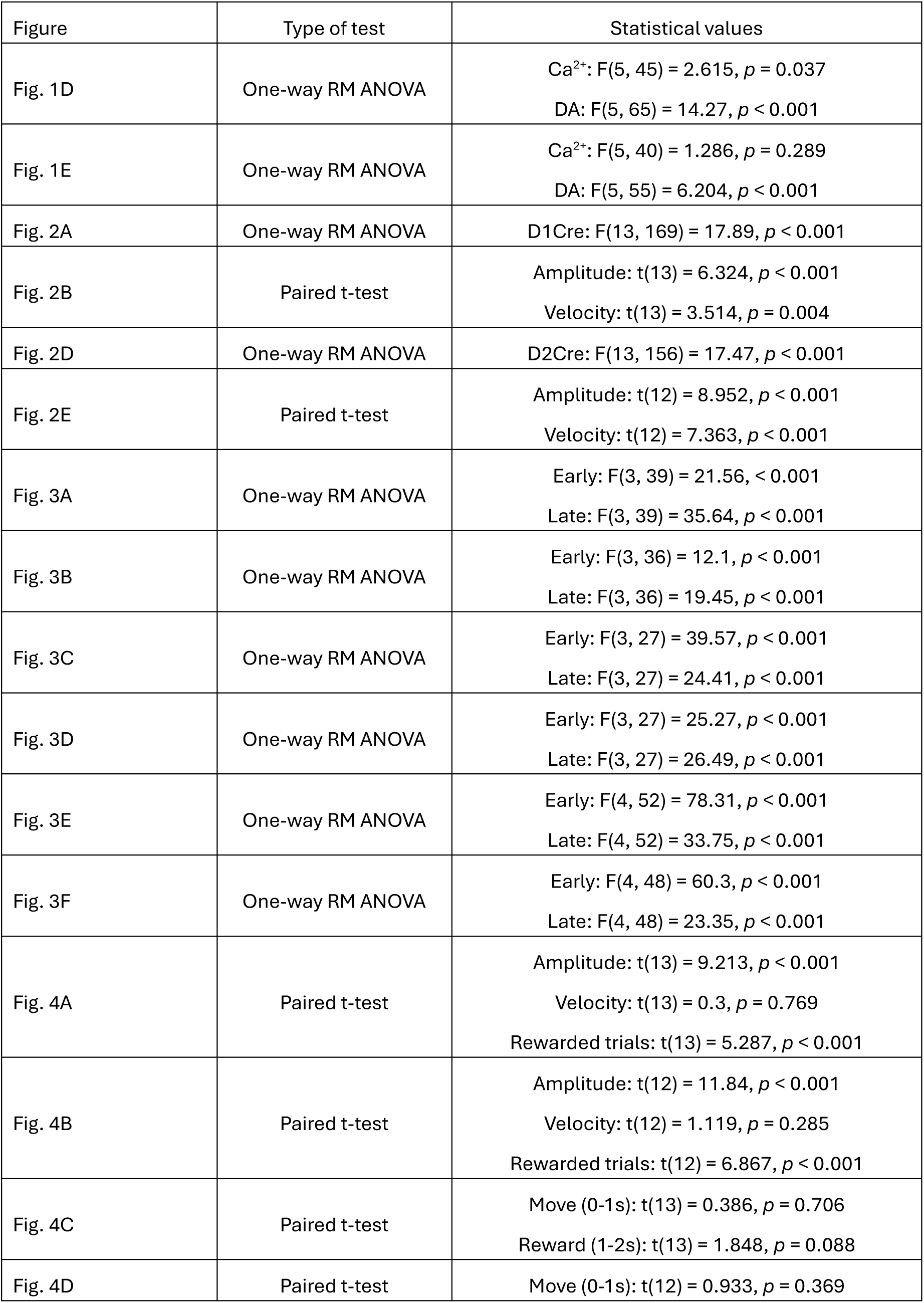
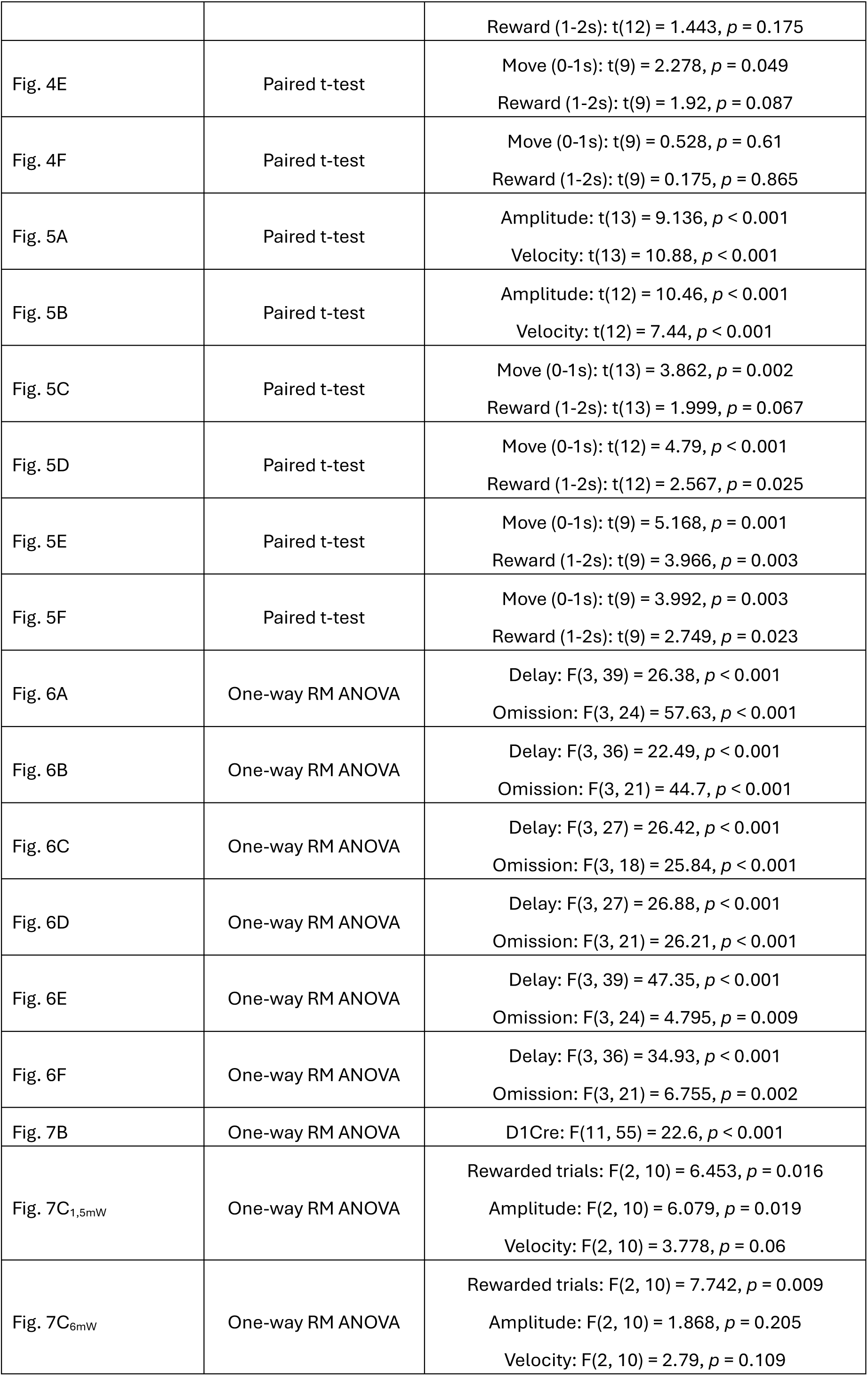
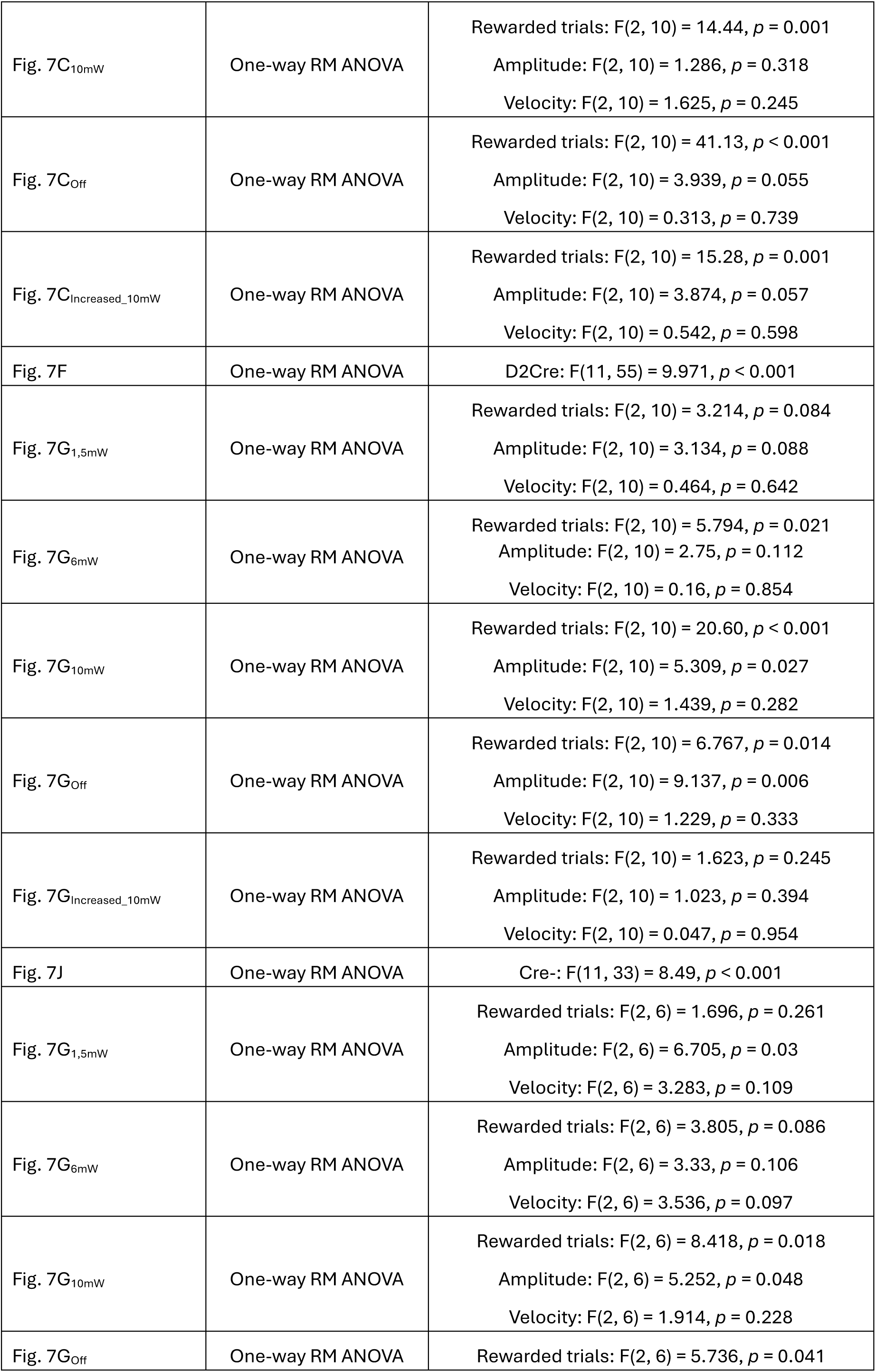
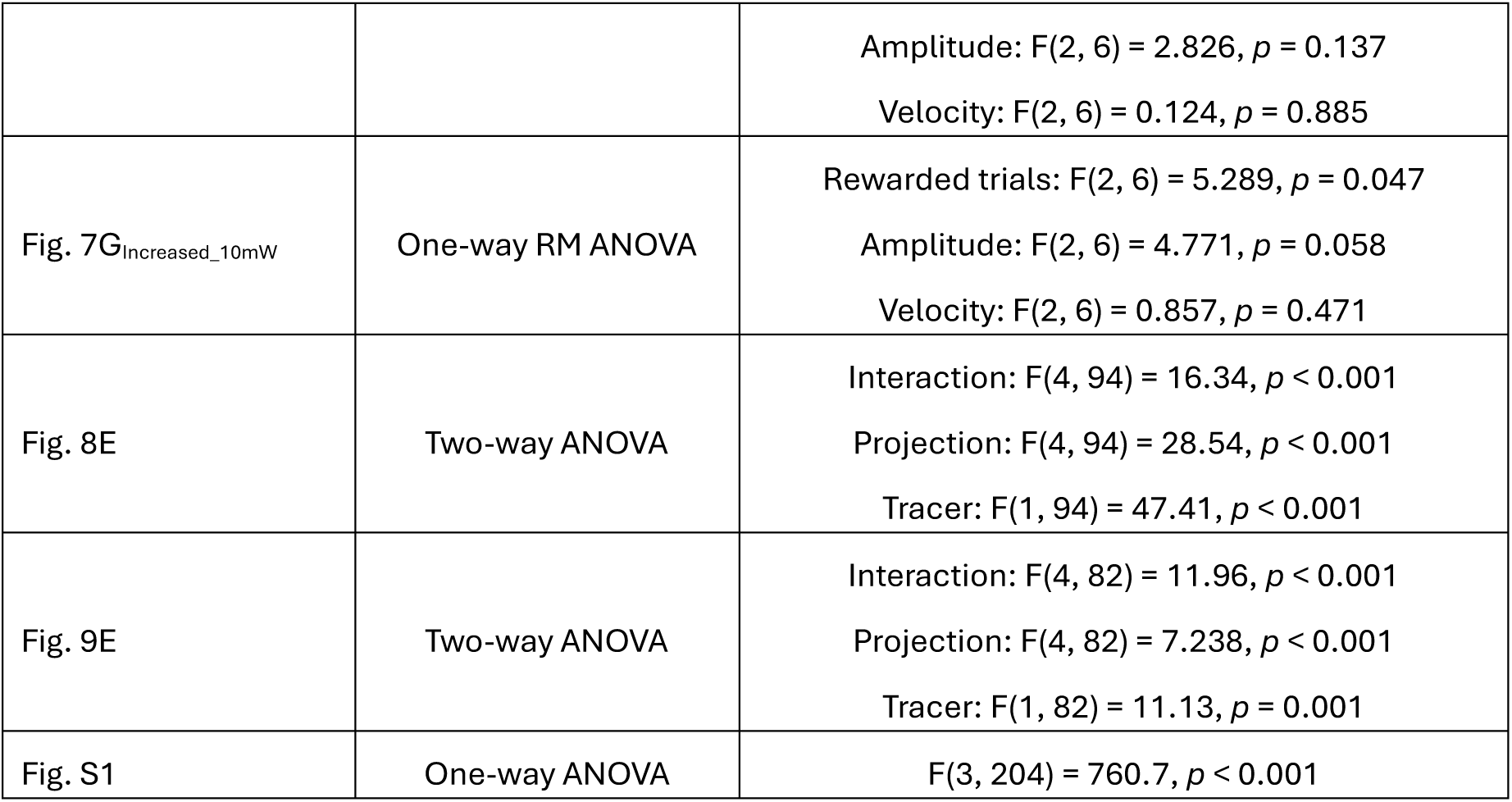
Statistical table.

| Figure | Type of test | Statistical values |
|---|---|---|
| Fig. 1D | One-way RM ANOVA | Ca<sup>2+</sup>: F(5, 45) = 2.615, *p* = 0.037<br>DA: F(5, 65) = 14.27, *p* < 0.001 |
| Fig. 1E | One-way RM ANOVA | Ca<sup>2+</sup>: F(5, 40) = 1.286, *p* = 0.289<br>DA: F(5, 55) = 6.204, *p* < 0.001 |
| Fig. 2A | One-way RM ANOVA | D1Cre: F(13, 169) = 17.89, *p* < 0.001 |
| Fig. 2B | Paired t-test | Amplitude: t(13) = 6.324, *p* < 0.001<br>Velocity: t(13) = 3.514, *p* = 0.004 |
| Fig. 2D | One-way RM ANOVA | D2Cre: F(13, 156) = 17.47, *p* < 0.001 |
| Fig. 2E | Paired t-test | Amplitude: t(12) = 8.952, *p* < 0.001<br>Velocity: t(12) = 7.363, *p* < 0.001 |
| Fig. 3A | One-way RM ANOVA | Early: F(3, 39) = 21.56, < 0.001<br>Late: F(3, 39) = 35.64, *p* < 0.001 |
| Fig. 3B | One-way RM ANOVA | Early: F(3, 36) = 12.1, *p* < 0.001<br>Late: F(3, 36) = 19.45, *p* < 0.001 |
| Fig. 3C | One-way RM ANOVA | Early: F(3, 27) = 39.57, *p* < 0.001<br>Late: F(3, 27) = 24.41, *p* < 0.001 |
| Fig. 3D | One-way RM ANOVA | Early: F(3, 27) = 25.27, *p* < 0.001<br>Late: F(3, 27) = 26.49, *p* < 0.001 |
| Fig. 3E | One-way RM ANOVA | Early: F(4, 52) = 78.31, *p* < 0.001<br>Late: F(4, 52) = 33.75, *p* < 0.001 |
| Fig. 3F | One-way RM ANOVA | Early: F(4, 48) = 60.3, *p* < 0.001<br>Late: F(4, 48) = 23.35, *p* < 0.001 |
| Fig. 4A | Paired t-test | Amplitude: t(13) = 9.213, *p* < 0.001<br>Velocity: t(13) = 0.3, *p* = 0.769<br>Rewarded trials: t(13) = 5.287, *p* < 0.001 |
| Fig. 4B | Paired t-test | Amplitude: t(12) = 11.84, *p* < 0.001<br>Velocity: t(12) = 1.119, *p* = 0.285<br>Rewarded trials: t(12) = 6.867, *p* < 0.001 |
| Fig. 4C | Paired t-test | Move (0-1s): t(13) = 0.386, *p* = 0.706<br>Reward (1-2s): t(13) = 1.848, *p* = 0.088 |
| Fig. 4D | Paired t-test | Move (0-1s): t(12) = 0.933, *p* = 0.369<br>Reward (1-2s): t(12) = 1.443, *p* = 0.175 |
| Fig. 4E | Paired t-test | Move (0-1s): t(9) = 2.278, *p* = 0.049<br>Reward (1-2s): t(9) = 1.92, *p* = 0.087 |
| Fig. 4F | Paired t-test | Move (0-1s): t(9) = 0.528, *p* = 0.61<br>Reward (1-2s): t(9) = 0.175, *p* = 0.865 |
| Fig. 5A | Paired t-test | Amplitude: t(13) = 9.136, *p* < 0.001<br>Velocity: t(13) = 10.88, *p* < 0.001 |
| Fig. 5B | Paired t-test | Amplitude: t(12) = 10.46, *p* < 0.001<br>Velocity: t(12) = 7.44, *p* < 0.001 |
| Fig. 5C | Paired t-test | Move (0-1s): t(13) = 3.862, *p* = 0.002<br>Reward (1-2s): t(13) = 1.999, *p* = 0.067 |
| Fig. 5D | Paired t-test | Move (0-1s): t(12) = 4.79, *p* < 0.001<br>Reward (1-2s): t(12) = 2.567, *p* = 0.025 |
| Fig. 5E | Paired t-test | Move (0-1s): t(9) = 5.168, *p* = 0.001<br>Reward (1-2s): t(9) = 3.966, *p* = 0.003 |
| Fig. 5F | Paired t-test | Move (0-1s): t(9) = 3.992, *p* = 0.003<br>Reward (1-2s): t(9) = 2.749, *p* = 0.023 |
| Fig. 6A | One-way RM ANOVA | Delay: F(3, 39) = 26.38, *p* < 0.001<br>Omission: F(3, 24) = 57.63, *p* < 0.001 |
| Fig. 6B | One-way RM ANOVA | Delay: F(3, 36) = 22.49, *p* < 0.001<br>Omission: F(3, 21) = 44.7, *p* < 0.001 |
| Fig. 6C | One-way RM ANOVA | Delay: F(3, 27) = 26.42, *p* < 0.001<br>Omission: F(3, 18) = 25.84, *p* < 0.001 |
| Fig. 6D | One-way RM ANOVA | Delay: F(3, 27) = 26.88, *p* < 0.001<br>Omission: F(3, 21) = 26.21, *p* < 0.001 |
| Fig. 6E | One-way RM ANOVA | Delay: F(3, 39) = 47.35, *p* < 0.001<br>Omission: F(3, 24) = 4.795, *p* = 0.009 |
| Fig. 6F | One-way RM ANOVA | Delay: F(3, 36) = 34.93, *p* < 0.001<br>Omission: F(3, 21) = 6.755, *p* = 0.002 |
| Fig. 7B | One-way RM ANOVA | D1Cre: F(11, 55) = 22.6, *p* < 0.001 |
| Fig. 7C<sub>1,5mW</sub> | One-way RM ANOVA | Rewarded trials: F(2, 10) = 6.453, *p* = 0.016<br>Amplitude: F(2, 10) = 6.079, *p* = 0.019<br>Velocity: F(2, 10) = 3.778, *p* = 0.06 |
| Fig. 7C<sub>6mW</sub> | One-way RM ANOVA | Rewarded trials: F(2, 10) = 7.742, *p* = 0.009<br>Amplitude: F(2, 10) = 1.868, *p* = 0.205<br>Velocity: F(2, 10) = 2.79, *p* = 0.109 |
| Fig. 7C<sub>10mW</sub> | One-way RM ANOVA | Rewarded trials: F(2, 10) = 14.44, *p* = 0.001<br>Amplitude: F(2, 10) = 1.286, *p* = 0.318<br>Velocity: F(2, 10) = 1.625, *p* = 0.245 |
| Fig. 7C<sub>Off</sub> | One-way RM ANOVA | Rewarded trials: F(2, 10) = 41.13, *p* < 0.001<br>Amplitude: F(2, 10) = 3.939, *p* = 0.055<br>Velocity: F(2, 10) = 0.313, *p* = 0.739 |
| Fig. 7C<sub>Increased_10mW</sub> | One-way RM ANOVA | Rewarded trials: F(2, 10) = 15.28, *p* = 0.001<br>Amplitude: F(2, 10) = 3.874, *p* = 0.057<br>Velocity: F(2, 10) = 0.542, *p* = 0.598 |
| Fig. 7F | One-way RM ANOVA | D2Cre: F(11, 55) = 9.971, *p* < 0.001 |
| Fig. 7G<sub>1,5mW</sub> | One-way RM ANOVA | Rewarded trials: F(2, 10) = 3.214, *p* = 0.084<br>Amplitude: F(2, 10) = 3.134, *p* = 0.088<br>Velocity: F(2, 10) = 0.464, *p* = 0.642 |
| Fig. 7G<sub>6mW</sub> | One-way RM ANOVA | Rewarded trials: F(2, 10) = 5.794, *p* = 0.021<br>Amplitude: F(2, 10) = 2.75, *p* = 0.112<br>Velocity: F(2, 10) = 0.16, *p* = 0.854 |
| Fig. 7G<sub>10mW</sub> | One-way RM ANOVA | Rewarded trials: F(2, 10) = 20.60, *p* < 0.001<br>Amplitude: F(2, 10) = 5.309, *p* = 0.027<br>Velocity: F(2, 10) = 1.439, *p* = 0.282 |
| Fig. 7G<sub>Off</sub> | One-way RM ANOVA | Rewarded trials: F(2, 10) = 6.767, *p* = 0.014<br>Amplitude: F(2, 10) = 9.137, *p* = 0.006<br>Velocity: F(2, 10) = 1.229, *p* = 0.333 |
| Fig. 7G<sub>Increased_10mW</sub> | One-way RM ANOVA | Rewarded trials: F(2, 10) = 1.623, *p* = 0.245<br>Amplitude: F(2, 10) = 1.023, *p* = 0.394<br>Velocity: F(2, 10) = 0.047, *p* = 0.954 |
| Fig. 7J | One-way RM ANOVA | Cre-: F(11, 33) = 8.49, *p* < 0.001 |
| Fig. 7G<sub>1,5mW</sub> | One-way RM ANOVA | Rewarded trials: F(2, 6) = 1.696, *p* = 0.261<br>Amplitude: F(2, 6) = 6.705, *p* = 0.03<br>Velocity: F(2, 6) = 3.283, *p* = 0.109 |
| Fig. 7G<sub>6mW</sub> | One-way RM ANOVA | Rewarded trials: F(2, 6) = 3.805, *p* = 0.086<br>Amplitude: F(2, 6) = 3.33, *p* = 0.106<br>Velocity: F(2, 6) = 3.536, *p* = 0.097 |
| Fig. 7G<sub>10mW</sub> | One-way RM ANOVA | Rewarded trials: F(2, 6) = 8.418, *p* = 0.018<br>Amplitude: F(2, 6) = 5.252, *p* = 0.048<br>Velocity: F(2, 6) = 1.914, *p* = 0.228 |
| Fig. 7G<sub>Off</sub> | One-way RM ANOVA | Rewarded trials: F(2, 6) = 5.736, *p* = 0.041<br>Amplitude: F(2, 6) = 2.826, *p* = 0.137<br>Velocity: F(2, 6) = 0.124, *p* = 0.885 |
| Fig. 7G<sub>Increased_10mW</sub> | One-way RM ANOVA | Rewarded trials: F(2, 6) = 5.289, *p* = 0.047<br>Amplitude: F(2, 6) = 4.771, *p* = 0.058<br>Velocity: F(2, 6) = 0.857, *p* = 0.471 |
| Fig. 8E | Two-way ANOVA | Interaction: F(4, 94) = 16.34, *p* < 0.001<br>Projection: F(4, 94) = 28.54, *p* < 0.001<br>Tracer: F(1, 94) = 47.41, *p* < 0.001 |
| Fig. 9E | Two-way ANOVA | Interaction: F(4, 82) = 11.96, *p* < 0.001<br>Projection: F(4, 82) = 7.238, *p* < 0.001<br>Tracer: F(1, 82) = 11.13, *p* = 0.001 |
| Fig. S1 | One-way ANOVA | F(3, 204) = 760.7, *p* < 0.001 |

### Key resources table

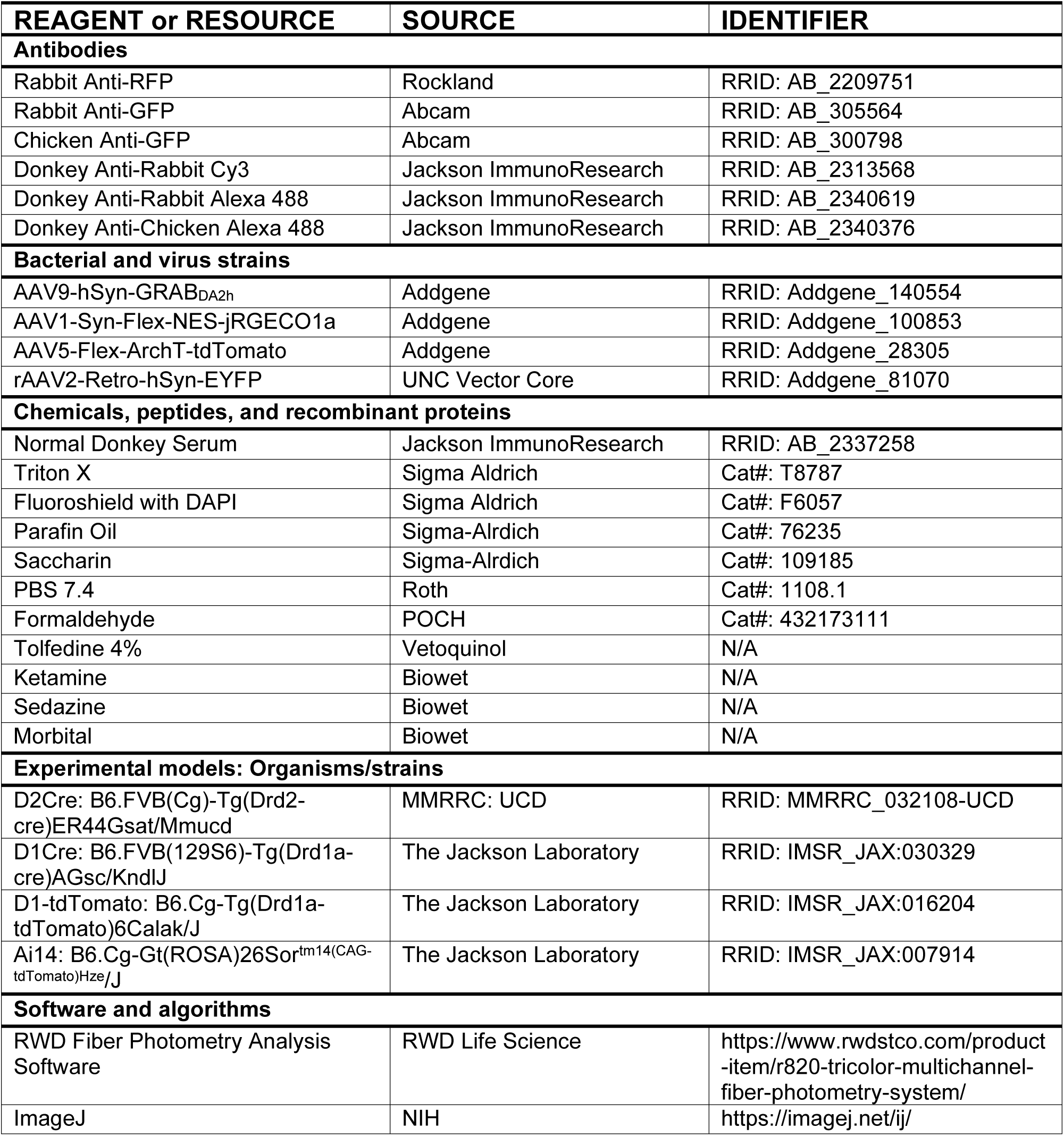
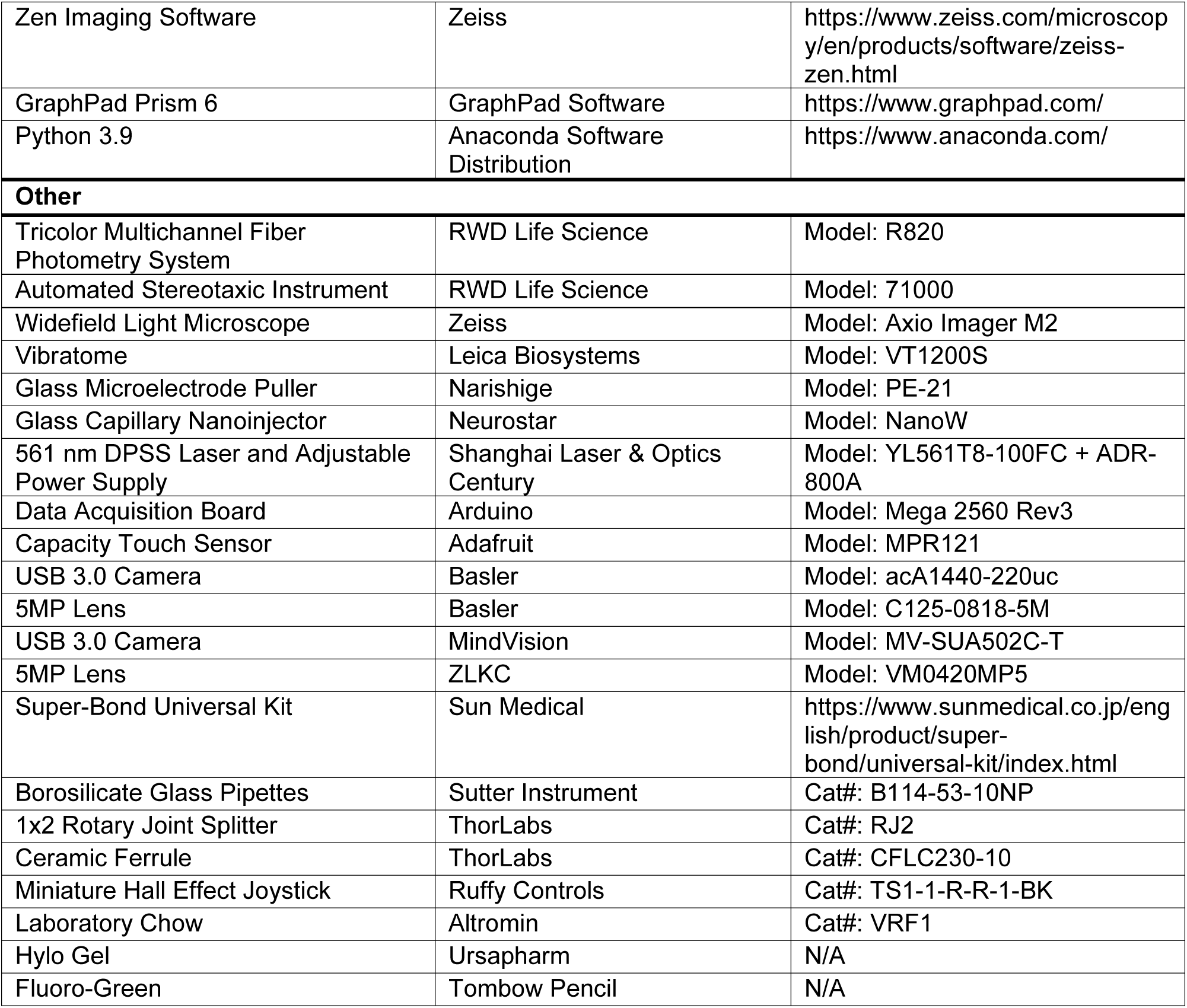

