## Supplemental information for "The role of dopamine-sensitive motor cortical circuits in the development and execution of skilled forelimb movements"

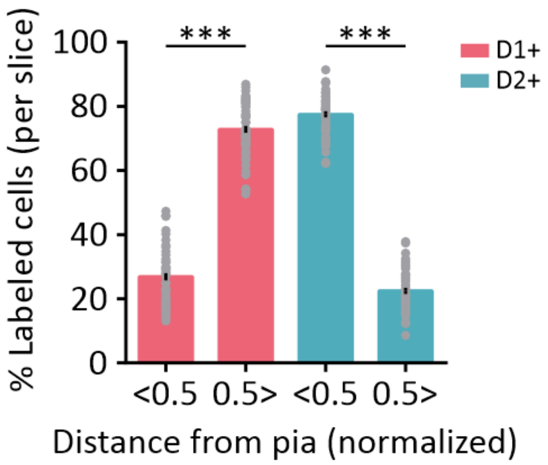


**Figure S1. D1+ and D2+ cells are distinctly distributed throughout layers of the M1.**

Percentage cell density of D1+ and D2+ neurons as a function of distance from pia (<0.5 superficial layers; 0.5> deep layers) (D1Cre: n = 11 mice, n = 56 slices; D2Cre: n = 10 mice, n = 48 slices). Data are represented as mean ± SEM. One-way ANOVA; multiple comparisons: each mean vs every other mean; Bonferroni's post hoc test; ****p* < 0.001.

**Video S1. D1Cre_jRGECO1a_S3.**

**Video S2. D1Cre_GRAB$_{DA2h}$_S3.**

**Video S3. D1Cre_jRGECO1a_S9.**

**Video S4. D1Cre_GRAB$_{DA2h}$_S9.**

**Video S5. D2Cre_jRGECO1a_S3.**

**Video S6. D2Cre_GRAB$_{DA2h}$_S3.**

**Video S7. D2Cre_jRGECO1a_S9.**

**Video S8. D2Cre_GRAB$_{DA2h}$_S9.**
